# Directing egg traffic: Internal mechanosensory feedback modulates rhythmic motor activity to coordinate ovulation in *Drosophila*

**DOI:** 10.1101/2025.09.22.677773

**Authors:** Shao-Cong Su, Ning Zhang, Chen-Yu Li, Ji-Yang Xing, Dick R. Nässel, Cong-Fen Gao, Shun-Fan Wu

## Abstract

Ovulation is a prerequisite for successful reproduction and requires precise coordination of muscle contractions, especially to propel eggs from the paired lateral oviducts into the common oviduct. Although internal mechanosensation is expected to provide critical feedback for this process, the sensorimotor circuitry and mechanisms by which sensory feedback shapes motoneuron activity and coordinates oviduct contractions are still poorly known. Here, in *Drosophila,* we identify a novel pair of multidendritic mechanosensory neurons (mdn-LO) in the lateral oviducts that express the mechanosensitive channels TMC (transmembrane channel-like protein), PPK (pickpocket), and Piezo. We show that TMC is essential for coordinated ovulation; *tmc* mutation or neuron-specific knockdown increases egg-jamming at the junction between the lateral and the common oviduct. In contrast, *ppk* or *Piezo* mutants display reduced egg-laying but no jamming. Calcium imaging reveals that the mdn-LO neurons are rhythmically activated by oviduct contractions, and chemogenetic stimulation of these neurons triggers muscle contractions. Genetic manipulations of the mdn-LO neurons disrupt egg passage and induce egg-jamming. The axons of the mdn-LO neurons form functional contacts with female-specific oviduct motoneurons in the abdominal neuromeres that express insulin-like peptide 7 (ILP7) and glutamate. Activation of mdn-LO neurons triggers Ca²⁺ activity in ILP7 motoneurons and elicits rhythmic oviduct contractions. Importantly, the ILP7 motoneurons exhibit spontaneous rhythmic Ca²⁺ activity that is modulated by oviduct contractions and egg movement. Furthermore, triggering activity in the ILP7 neurons induces egg-jamming. Finally, knockdown of ILP7 peptide affects the rate of ovulation, but plays no role in egg jamming, which instead is likely induced by colocalized glutamate. In summary, our study identifies a novel circuit, including the TMC-expressing mdn-LO neurons and a small set of ILP7 motoneurons, that organizes an innate behavior by coupling strategic sensory information and rhythmical motor output to ensure bilateral coordination of oviduct contractions to prevent egg jamming during oviposition.

**Significance:** Ovulation requires precisely coordinated contractions of the lateral oviducts to propel eggs into the common oviduct without jamming. Yet the sensory cells and mechanotransduction channels that coordinate movement through the paired lateral oviducts remain unclear. Here, we identify a novel pair of lateral-oviduct multidendritic neurons (mdn-LO) that express the conserved mechanosensory channels TMC (transmembrane channel-like protein), PPK (pickpocket), and Piezo, and show that TMC is uniquely required to prevent egg jamming at the junction between the lateral and common oviducts. The mdn-LO neurons are activated by oviduct contractions and relay signals to female-specific glutamatergic motoneurons in the abdominal neuromere. These paired oviduct motoneurons are rhythmically active and regulate contractions of the lateral oviducts. Together, our findings demonstrate that TMC-mediated mechanosensory feedback fine-tunes motor output to ensure coordinated ovulation and successful reproduction.

## Introduction

Successful reproduction in oviparous organisms relies on the precise coordination of muscle contractions during ovulation and oviposition, a process orchestrated by neural circuits that generate rhythmic motor outputs and sensory feedback. Studies of the genetically tractable insect *Drosophila* have shown that successful reproduction requires strict and precise regulation of events to occur in coordination, including mechanisms and behaviors associated with oogenesis, follicle cell rupture, motor control of reproductive oviduct muscles to ensure ovulation, egg activation, the choice of egg-laying site, and finally egg deposit<u>1</u>, <u>2</u>, <u>3</u>, <u>4</u>, <u>5</u>, <u>6</u>. In *Drosophila,* the neuronal circuitry involved in mating and post-mating is well understood <u>7</u>, <u>8</u>, <u>9</u>, <u>10</u>, <u>11</u>, <u>12</u>, <u>13</u>, <u>14</u>, <u>15</u>, <u>16</u>, and several aspects of the motor control of ovulation and oviposition have been studied <u>4</u>, <u>8</u>, <u>17</u>, <u>18</u>. One critical obstacle during ovulation is the passage of eggs from the paired ovaries into the common oviduct. To avoid jamming of eggs at the junction between the two lateral oviducts, critical mechanosensory feedback is required to ensure sequential activity in motoneurons to generate alternating contractions of left and right oviduct muscle. The internal sensory neurons providing feedback and motor circuitry responsible for this directing of egg traffic into the common oviduct have not been identified. Here, we untangle the neuronal components responsible for this coordinated egg transport, starting with sensory neurons in the oviduct and proceeding to motoneurons in the abdominal neuromeres of the ventral nerve cord (VNC).

Previous studies have identified some sensory neurons in the oviduct. The contractions of the common oviduct can be sensed by mechanosensory neurons in the common oviduct that express pickpocket (PPK) <u>8</u>. Further down the common oviduct, a pair of posterior uterine sensory neurons (PU neurons) sense the passage of eggs into the ovipositor <u>1</u>. However, it remains unclear which neurons are responsible for sensing the contractions of the lateral oviducts. Here, we identify a novel pair of multidendritic mechanosensory neurons in the lateral oviducts (designated mdn-LO) that express three evolutionarily conserved mechanosensitive channels: TMC (transmembrane channel-like protein), PPK (pickpocket), and Piezo, along with the sex-determination gene *doublesex* (distinct from the adjacent *fruitless*-expressing multidendritic neurons). We show that TMC is uniquely required for ovulation coordination, in contrast to PPK and Piezo. TMC mutation or neuron-specific knockdown causes severe egg-jamming. Calcium imaging confirms that mdn-LO neurons are rhythmically activated by lateral oviduct contractions, consistent with a mechanosensory role.

What about motoneurons regulating contractions of the lateral oviduct muscle? Two types of efferent neurons that supply axons to the oviduct muscle have been identified. The first is a set of efferent octopaminergic neurons identified in the abdominal neuromeres of the VNC. These release octopamine (OA) that acts on two different receptors (OAMB and Octβ2R) to cause a series of contractions and relaxations of the muscle in the oviducts to move the egg forward <u>17</u>, <u>18</u>, <u>19</u>, <u>20</u>, <u>21</u>. More specifically, OA acts on the OAMB receptor to allow entrance of the egg into one of the lateral oviducts by the rupture of the posterior mature follicle cells surrounding the oocyte <u>5</u>. Further movement through the lateral and common oviducts is caused by OA modulation of contractions in the myogenic muscle in the reproductive tract <u>17</u>, <u>18</u>, <u>19</u>, <u>20</u>.

In addition to OA regulation of oviduct contractions, a group of female-specific glutamatergic motor neurons located in the abdominal neuromeres has been identified. These bilateral pairs of neurons co-express insulin-like peptide 7 (ILP7) and were shown to induce contractions of the lateral and the common oviducts <u>4</u>, <u>8</u>, <u>22</u>. In the following, we call these neurons ILP7-neurons. We show here that silencing of these ILP7-neurons causes jamming of eggs at the junction between the common and lateral oviducts. Using trans-Tango and GRASP analyses together with calcium imaging, we demonstrate that mdn-LO neurons form functional connections with the ILP7 neurons in the abdominal neuromeres. Activation of either mdn-LO or ILP7 neurons induces egg jamming. Moreover, ILP7 neurons exhibit spontaneous rhythmic calcium oscillations that are modulated by mechanosensory input from the lateral oviducts. Thus, acting in concert, the left and right mechanosensory mdn-LO cells and ILP7 motoneurons generate the alternating contractions in the lateral oviducts necessary for coordinated egg entrance into the common oviduct. Although ILP7 neuropeptide is produced by the ILP7-neurons, knockdown of the peptide does not affect egg jamming, but reduces the number of eggs laid, suggesting that its action may be on the common oviduct rather than the lateral ones. In summary, we outline a circuit consisting of a pair of TMC-dependent mechanosensory neurons and a set of rhythmically active ILP7 oviduct motoneurons that ensures smooth passage of eggs through the lateral oviducts and thereby successful oviposition and reproduction. We furthermore describe a novel function of the evolutionarily conserved TMC channels.

## Results

### Loss of the transmembrane channel-like (tmc) gene leads to egg-jamming in Drosophila

The paired oviducts connect the ovaries to the uterus and are essential for successful egg transport during ovulation. In *Drosophila*, typically only one egg traverses the oviduct at a time, indicating that internal sensory mechanisms coordinate egg movement to prevent egg jamming. Regarding internal sensation within the female reproductive system, existing reports are limited to *ppk*-expressing neurons in the common oviduct that enhance *Drosophila*’s preference for acetic acid by sensing oviduct contractions <u>8</u>, as well as posterior uterine (PU) sensory neurons that are activated by egg progression <u>1</u>. However, how internal sensory mechanisms coordinate the process of egg transport from the two lateral into the unpaired uterus to avoid egg jamming remains unclear.

In addition to Pickpocket (PPK), PIEZO, and Transmembrane Channel-like (TMC) proteins have also been identified in the female reproductive system of *Drosophila*, though their functional roles remain unclear <u>8</u>, <u>23</u>. PPK, PIEZO, and TMC are evolutionarily conserved mechanosensitive ion channels <u>8</u>, <u>23</u>, <u>24</u>, <u>25</u>, <u>26</u>, <u>27</u>, <u>28</u>. In *Drosophila* these channels have been implicated in the regulation of sexual behavior <u>29</u>, <u>30</u>, oviposition preference <u>28</u>, and feeding behavior <u>23</u>, <u>31</u>.

Given our previous finding that *Nltmc3*, a TMC ortholog in the brown planthopper (BPH) *Nilaparvata lugens*, is essential for normal ovary development and oviposition<u>32</u>, we sought to investigate whether the *Drosophila* ortholog *tmc* (CG46121) plays a similar role. Using *tmc* mutant flies, *tmc^1^* <u>23</u>, we conducted oviposition assays and found that egg-laying was significantly reduced compared to controls (Fig. 1A).

**Figure 1.**
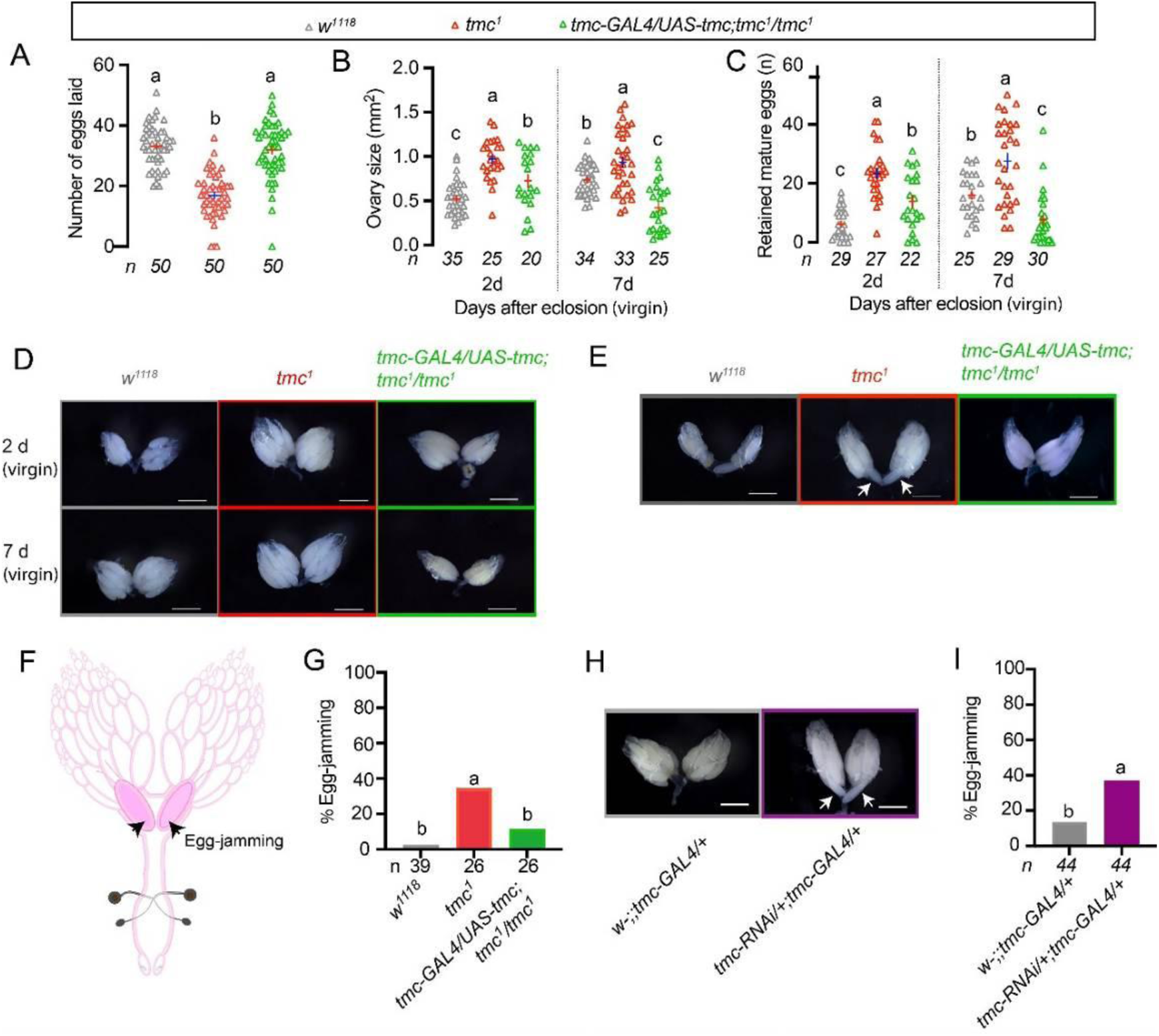
The *Tmc* channel is essential for ovulation and oviposition in *Drosophila*. *Tmc* mutants lay fewer eggs and display ovulation defects. All experiments, except those in Fig. 1B-D, used mated flies. Groups that share at least one letter are not statistically distinguishable (*P* > 0.05), whereas groups with different letters are statistically distinguishable (*P* < 0.05). (**A**) The number of eggs laid per female of the *w^1118^*, *tmc^1^* mutant, and *tmc-GAL4/UAS-tmc*; *tmc^1^* (mutant with *tmc* selectively rescued in *tmc*-GAL4-expressing cells) for 14 h at 25 °C. One-way ANOVA with Tukey’s Multiple Range Test. (**B-C**) Quantification was performed on the ovary size and the number of mature eggs retained in each pair of ovaries from *w^1118^*, *tmc^1^*, and *tmc-GAL4/UAS-tmc;tmc^1^* flies. The results show that unmated *tmc^1^* mutant females retained more mature eggs in their ovaries, resulting in larger ovaries. This phenotype can be rescued by expressing *tmc* in neurons specifically marked by *tmc-GAL4*. Representative images of the ovaries are shown in Figure 1D. One-way ANOVA with Tukey’s Multiple Range Test. (**D**) Representative images of *w^1118^*, *tmc^1^*, *tmc-GAL4/UAS-tmc*; *tmc^1^* ovaries on d2AAE (2 d after eclosion) and d7AEE (7 d after eclosion), from unmated flies. (**E**) Representative images of *w^1118^*, *tmc^1^*, *tmc-GAL4/UAS-tmc*; *tmc^1^*/*tmc^1^* ovaries. White arrows mark the egg in the lateral oviduct. (**F**) Illustration of egg-jamming in the lateral oviduct. (**G**) The egg-jamming ratio (in %). Genotypes as in Fig 1E. Fisher’s exact test. (**H**) Representative images of *w-;;tmc-GAL4/+, tmc-RNAi/+;tmc-GAL4/+* ovaries. White arrows mark the egg in the lateral oviduct. (I) The egg-jamming ratio (in %). Genotypes as in Fig. 1H. Fisher’s exact test.

**Figure 1 — figure supplement 1.**
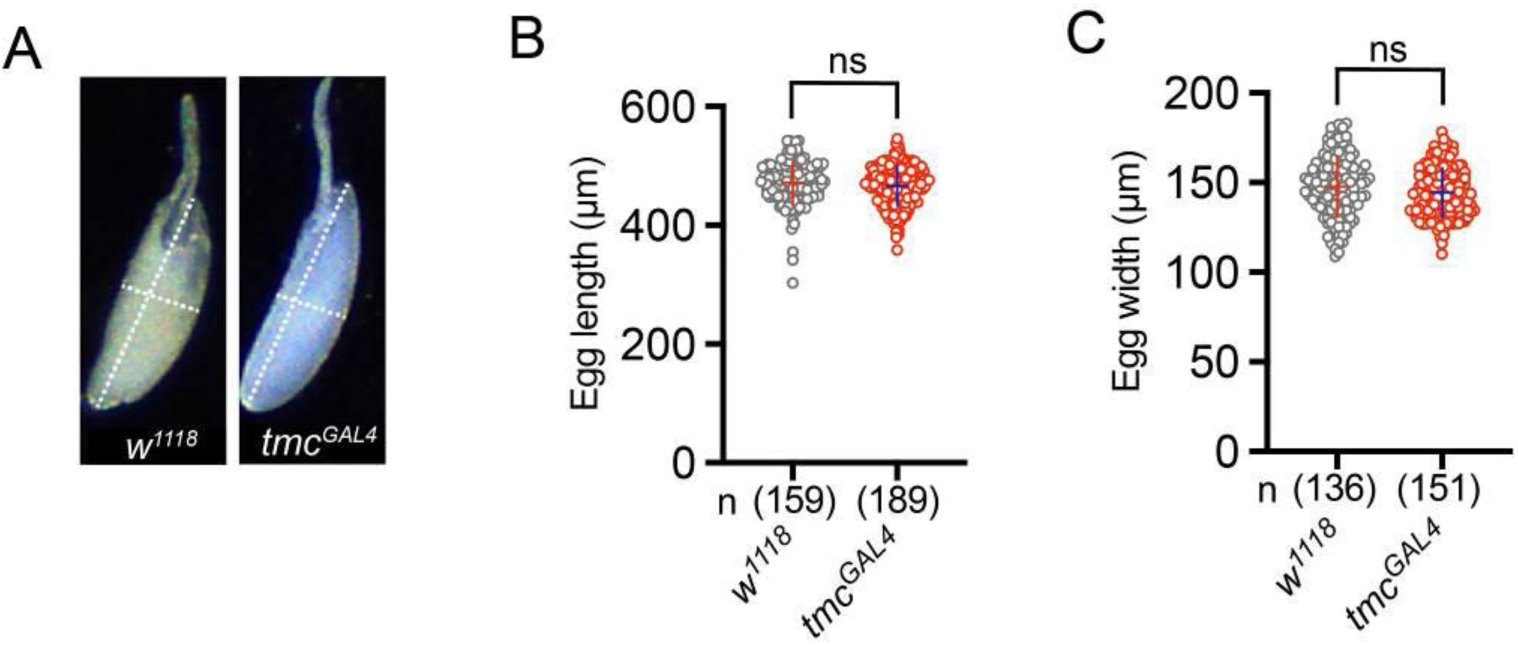
The *tmc* gene does not affect egg shape. (**A**) Representative images of eggs of *w^1118^* flies and the *tmc^GAL4^* mutant. (**B – C**) The egg length and width of the *w^1118^* and *tmc^GAL4^* mutants. Groups that share at least one letter are not statistically distinguishable (*P* > 0.05), whereas groups with completely different letters are statistically distinguishable (*P* < 0.05); Student’s t-test.

**Figure 1 – figure supplement 2.**
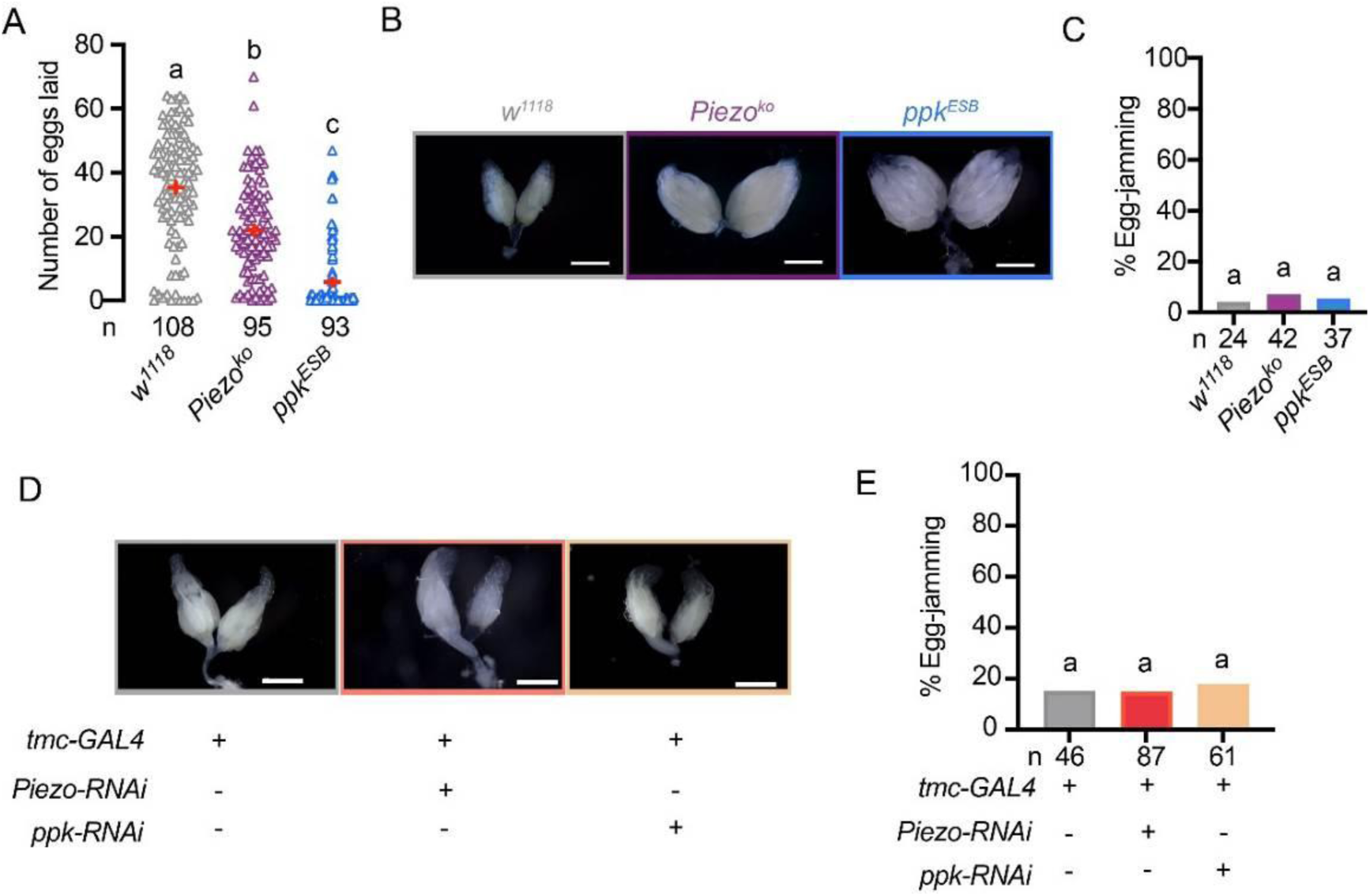
Egg jamming is not affected by the manipulation of other mechanosensory channel genes. (**A**) Number of eggs laid per female of *w^1118^*, *Piezo^ko^, ppk^ESB^* flies for 14 h at 25 °C. One-way ANOVA with Tukey’s Multiple Range Test. In this and the panels **C and E**: groups that share at least one letter are not statistically distinguishable (*P* > 0.05), whereas groups with completely different letters are statistically distinguishable (*P* < 0.05). (**B**) Representative images of *w^1118^*, *Piezo^ko^*, *ppk^ESB^* ovaries. (**C**) The egg-jamming ratio. Genotypes as in **B**. Fisher’s exact test.(**D**) Representative images of *w-;;tmc-GAL4/+, ppk-RNAi/+; tmc-GAL4/+, Piezo-RNAi/+; tmc-GAL4/+* ovaries. (**E**) The egg-jamming ratio. Genotypes as in **D**. Fisher’s exact test.

Since reduced egg-laying can result either from defects in ovarian development or disruptions in ovulation, we first examined whether developmental effects contributed to the phenotype. To rule out effects from mating, we initially used virgin female flies (for other experiments, unless noted otherwise, mated females were used). We assessed ovarian development in virgin *tmc^1^* mutants by measuring ovary size and found no developmental abnormalities, in contrast to BPHs <u>32</u>. Instead, we observed significantly enlarged ovaries in *tmc^1^* flies at both day 2 and day 7 post-eclosion compared to *w^1118^* controls (Fig. 1B, D). The enlarged ovaries were due to the accumulation of mature eggs, suggesting a severe ovulation defect in these unmated flies (Fig. 1C, D), which was not attributable to abnormalities in egg morphology (Fig. 1—figure supplement 1).

To further confirm the role of TMC in ovulation, we reintroduced the *tmc* gene into *tmc^1^* mutants using the *tmc-GAL4* driver line <u>23</u>. The reduced oviposition, enlarged ovaries, and accumulation of mature eggs observed in *tmc^1^* mutants were all rescued by expressing *UAS-tmc* under the *tmc* promoter (Fig. 1A–D).

We next dissected mated *tmc* mutant females to assess ovulation. Interestingly, we occasionally observed egg-jamming at the junction where the two lateral oviducts merge into the common oviduct (Fig. 1E and Fig. 1F, respectively). The frequency of this blockage was significantly higher in *tmc* mutants than in controls. Importantly, the reintroduction of the *tmc* gene in *tmc^1^* mutants eliminated this egg-jamming phenotype (Fig.1G). Similarly, RNAi-mediated knockdown of *tmc* in *tmc-GAL4*-expressing cells also resulted in egg-jamming (Fig. 1H–I).

Finally, we explored whether the other two channels might contribute to oviposition and ovulation regulation in *Drosophila* (Figure 1—figure supplement 2). Using *ppk* mutants (*ppk^ESB^*) and *Piezo* mutants (*Piezo^KO^*), we observed a significant reduction in egg-laying in both strains after mating (Figure 1—figure supplement 2A). However, we did not observe increased egg-jamming in these mutants or in flies with RNAi-mediated knockdown of *ppk* or *Piezo* in TMC-GAL4-expressing cells (Figure 1—figure supplement 2B–E).

These findings show that TMC, specifically in *tmc-GAL4*-expressing cells, plays a key role in coordinating normal ovulation in the lateral oviducts of *Drosophila*, whereas *Piezo* and PPK do not. The disruption of TMC leads to a significantly elevated incidence of egg-jamming, whereas the other channels may affect ovulation/oviposition via the common oviduct or uterus. The roles of Piezo and PPK in *tmc*-positive sensory neurons remain unresolved, underscoring the need for future studies to reveal the distinct contributions of these two mechanosensory channels and how the three channels together affect oviduct function.

### *Tmc* is localized in multi-dendritic neurons in the lateral oviduct

Whereas PPK and PIEZO have been previously localized to sensory neurons of the lateral oviducts <u>8</u>, the presence of TMC in the reproductive system was hitherto not known. Thus, we set out to examine the detailed expression of this channel using a *tmc-GAL4* and by immunohistochemistry. For the latter, we generated a TMC antiserum using a previously reported method <u>26</u>. As revealed by immunocytochemical labeling and *tmc-GAL4* expression, TMC is localized in a pair of multi-dendritic neurons in the lateral oviducts that we designate **m**ulti-**d**endritic **n**eurons in the **L**ateral **O**viduct, mdn-LO (Figure 2A-B). Double-labeling experiments with TMC antibodies and *tmc-GAL4-*driven green fluorescent protein (GFP) confirmed the expression of the channel protein in these mdn-LO neurons (Figure 2A). We did not observe anti-TMC immunoreactivity in *tmc^1^* mutant animals, confirming the specificity of the TMC antibodies (Figure 2C). To acquire high-resolution images of the processes of the *tmc-GAL4* labeled neurons, we expressed a membrane-bound GFP (*UAS-mCD8::GFP*) under the control of *tmc-GAL4.* We found that the *tmc-GAL4* specifically identified one mdn-LO neuron in each of the two bilaterally symmetrical lateral oviducts (Figure 2D). TMC is expressed in the dendritic branches, suggesting a sensory role (Figure 2D). Each of these *tmc* neurons covers the native lateral oviduct (the one with the neuronal cell body) with its dendrites (Figure 2D). We found no *tmc*-positive sensory neuron processes in the common oviduct or the uterus (Figure 2D), where branches of PPK-positive neurons have been detected <u>8</u>, <u>33</u>. We also investigated the neuronal expression of two *tmc-GAL4* lines (insertions on the second and third chromosomes, respectively) and a *tmc-LexA* line in the central nervous system. Similar to previous studies <u>23</u>, we found no *tmc*-positive cell bodies in the abdominal neuromeres (ABD) of the VNC in adult females <u>34</u> (Figure 2-supplemental file 1A-C), but there are many *tmc*-expressing neuronal processes projecting from peripheral tissues into the brain and posterior ABD. This part of the ABD is a region dedicated to processing female reproductive sensory input from the periphery and the origin of motor output to muscles involved in ovulation and egg-laying behavior <u>34</u> (Figure 2-supplemental file 1A-C).

**Figure 2.**
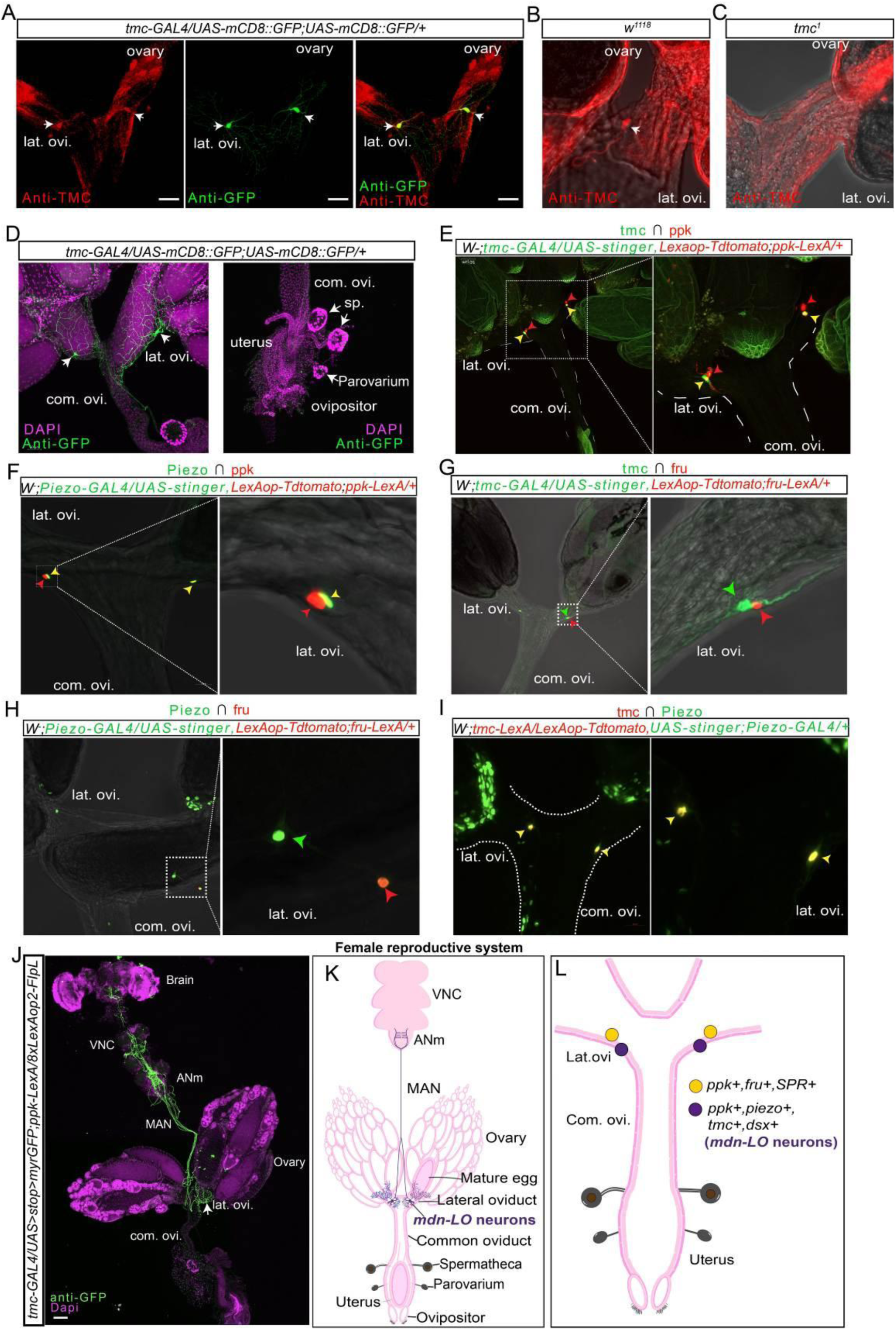
Expression pattern of mechanosensory channels in mdn-LO neurons of the lateral oviducts. **(A)** Labeling of a lateral oviduct (lat. ovi.) with antiserum to TMC and anti-GFP staining from flies expressing the *UAS-mCD8::GFP* reporter under the control of the *tmc-GAL4*. mdn-LO neurons are indicated by arrows. **(B – C)** Labeling of the lateral oviduct (lat. ovi.) using TMC antibodies in (B) *w^1118^* control and (C) *tmc^1^* mutant. The arrow points to one labeled mdn-LO neuron. **(D)** TMC is specifically expressed in a pair of multi-dendritic sensory neuron cells in the lateral oviduct. Even when the egg enters the lateral oviduct, the mdn-LO neuron dendrites are still wrapped around the lateral oviduct. *tmc* is not expressed in any other part of the female reproductive system: the common oviduct (com.ovi.), spermatheca (sp.) uterus, or parovaria (female accessory glands). **(E)** Lateral oviduct neurons (mdn-LO neurons) that express both *tmc* and *ppk* are revealed by a combinatorial strategy (indicated by yellow arrows). **(F)** Lateral oviduct neurons (mdn-LO neurons) that express both *Piezo* and *ppk* are revealed by a combinatorial strategy (indicated by yellow arrows). **(G)** Lateral oviduct neurons that express tmc (green arrows) and are labeled by the fruP1 driver (fruP1+) (red arrows). **(H)** Lateral oviduct neurons that express Piezo (green arrows) and are labeled by the fruP1 driver (fruP1+) (red arrows). **(I)** Lateral oviduct neurons(mdn-LO neurons) that express both *Piezo* and *ppk* are revealed by a combinatorial strategy (indicated by yellow arrows). **(J)** Expression of *tmc* and *ppk* expressing neurons in the brain, VNC, and lateral oviducts (lat. ovi.), using the intersectional technique (*GAL4/LexA*). The ANm region is targeted by mdn-LO neurons from the lateral oviducts. The genotype: *tmc-GAL4/UAS>stop>myrGFP;ppk-LexA/8xLexAop2-FIpL.* Common oviduct (com. ovi). median abdominal nerve (MAN). **(K)** Illustration of the morphology of the lateral oviduct *tmc-GAL4*-expressing neurons (mdn-LO) in the female reproductive system and ABD of the ventral nerve cord. **(L)** Summary illustration of the expression pattern of mechanosensory and other genes in lateral oviduct neurons. *Dsx*: doublesex.

**Figure 2—figure supplement 1.**
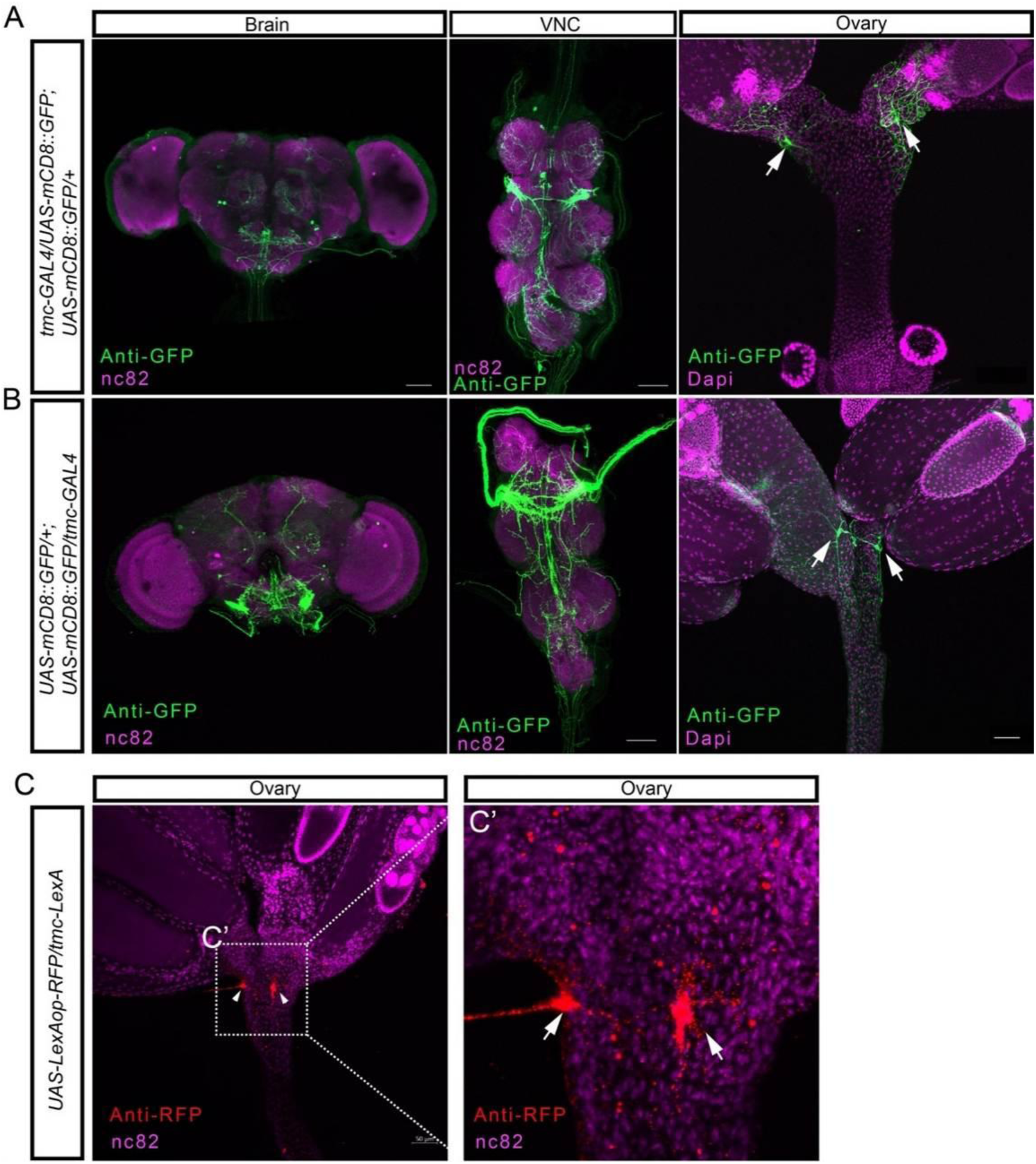
Expression patterns of different *tmc* drivers in the brain, ventral nerve cord, and reproductive system. (**A-B**) Expression patterns of *tmc-GAL4* (lines with insertion on different chromosomes) expressing neurons in the brain, ventral nerve cord (VNC), and ovary. (**C**) Expression of *tmc-LexA* expressing neurons (cell bodies indicated by white arrows) in the oviduct.

**Figure 2—figure supplement 2.**
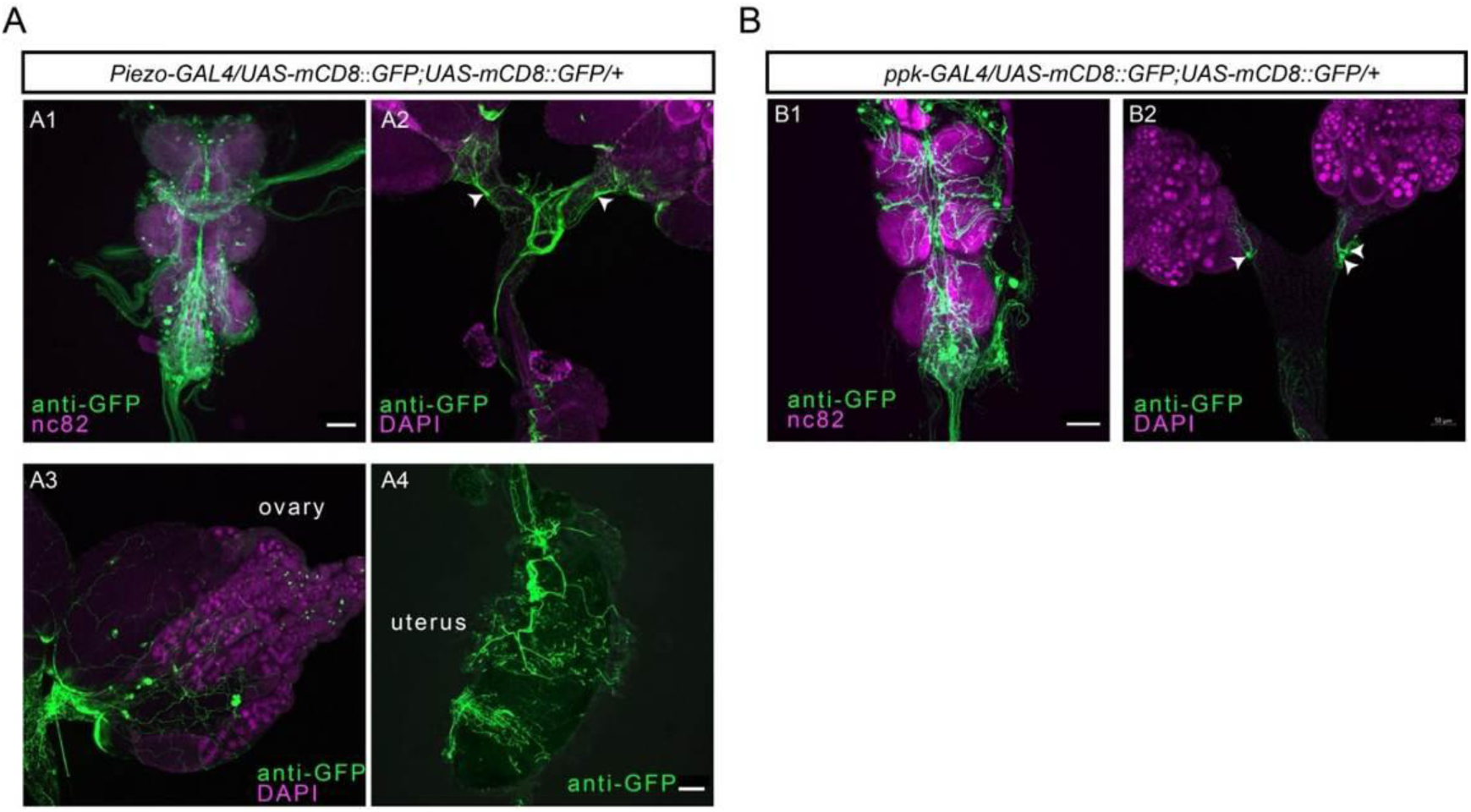
Expression of *Piezo* and *ppk* in CNS and reproductive system. (**A-B**) Expression patterns of *ppk* and *Piezo* in the female VNC (A1, B1) and reproductive system (**A2, B2**). (**A3, A4**) Expression patterns of *Piezo* in the ovary and uterus. White arrowheads mark the lateral oviduct neurons.

We next tested whether Piezo and PPK are expressed in neurons of the lateral oviduct. Similar to previous studies <u>8</u>, we observed both *Piezo-GAL4* and *PPK-GAL4* expression in multidendritic neurons in each lateral oviduct (Figure 2-supplemental file 2A and B). Then we asked whether TMC is co-expressed with PPK or PIEZO in neurons of the lateral oviduct. Indeed, double labeling of *PPK-LexA* and *tmc-GAL4* neurons revealed that the pair of TMC-expressing mdn-LO neurons is also PPK-positive (Figure 2E). We then employed an intersectional strategy <u>35</u>, <u>36</u> using a combination of *tmc-GAL4*, *PPK-LexA,* and FLP recombinase to label mdn-LO neurons and observed one pair of TMC and PPK co-expressing mdn-LO neurons (Figure 2J). The axons of this pair of neurons project from the reproductive system to the abdominal neuromeres of the VNC. (Figure 2J-K)

Using the same approach, we examined whether Piezo and PPK, TMC and Fruitless (fru), as well as TMC and Piezo, are co-expressed in neurons of the lateral oviduct (Figure 2F–I). Taken together, our results indicate that among the two pairs of multidendritic neurons located in the lateral oviducts, one pair (mdn-LO) expresses all three channels, TMC, PPK, and Piezo, as well as Dsx (Doublesex), while the other pair expresses only PPK and fru (Figure 2L) (for Dsx, see also Rezával et al., 2012). Note that FruM is male-specific and absent in females. Thus, the fru expression in females indicates fruP1 regulatory activity rather than FruM protein expression <u>37</u>.

### A pair of *Tmc*-expressing mdn-LO neurons in the lateral oviduct is essential for ovulation, and avoidance of egg-jamming

We selectively silenced the TMC-expressing neurons by using the *tmc-GAL4* to direct expression of the inward-rectifying potassium channel *Kir2.1* <u>38</u> and found that this manipulation causes animals to retain a similar number of eggs as the *tmc^1^*mutants do (Figure 3A). The binding of GAL80 to the C-terminal 30 amino acids of GAL4 can prevent GAL4-mediated transcriptional activation <u>39</u> and thus inhibit the activity of GAL4 in *Drosophila* <u>40</u>. We thus silenced *tmc*-*GAL4* neurons with *Kir2.1* and examined the effect of *elav-GAL80* suppression of the *tmc*-*GAL4*-mediated transgene expression in mdn-LO neurons. Interestingly, *tmc-GAL4 > Kir2.1* flies expressing *elav-GAL80* laid almost normal numbers of eggs compared to control flies, indicating that the oviposition phenotype arises predominantly from the *tmc*-GAL4 expressing mdn-LO neurons (Figure 3B). Taken together, these findings show that mdn-LO neurons are necessary for normal ovulation. Furthermore, we asked whether silencing *tmc*-expressing neurons also affects ovulation coordination in *Drosophila*. However, although acute silencing of *tmc-GAL4-*expressing mdn-LO neurons with Kir2.1 reduced oviposition, it did not increase the incidence of egg jamming at the junction between the lateral and common oviducts (Figure 3C-D). We interpret this difference from *tmc* loss-of-function mutants as follows: in *tmc* mutants, mdn-LO neurons are present and active but fail to properly transduce mechanical stimuli, thus producing a defective sensory drive to the motor circuit and thereby promoting jamming. In contrast, silencing mdn-LO removes the sensory drive altogether, yielding a state in which fewer eggs reach the junction, leading to reduced oviposition without increased jamming.

**Figure 3.**
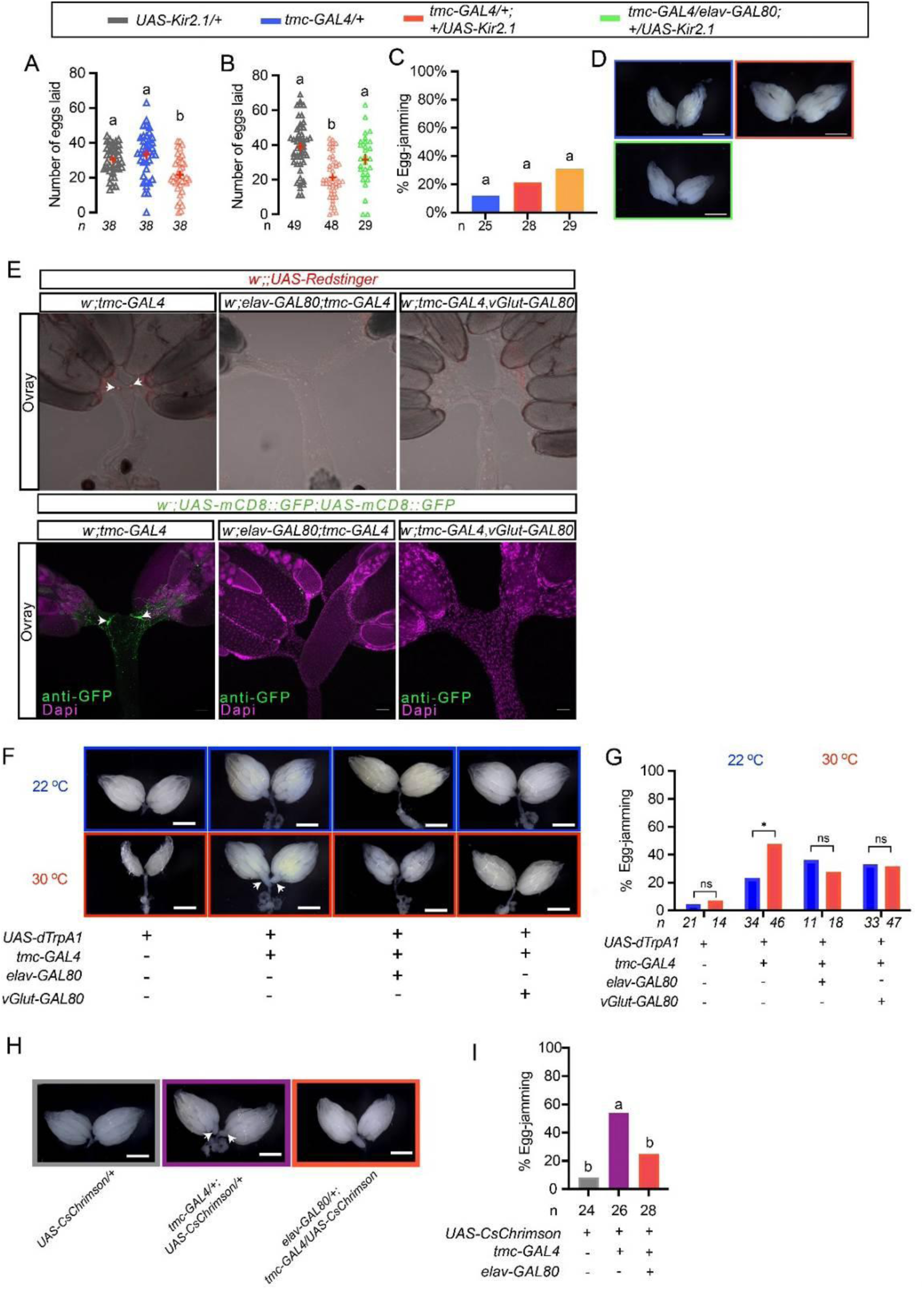
A pair of Tmc-expressing mdn-LO neurons in the lateral oviduct is essential for ovulation, and activation egg-jamming oviduct. In these graphs, \**P* < 0.05, \*\**P* < 0.01, \*\*\**P* < 0.001, \*\*\*\**P* < 0.0001, and ns (non-significant). Groups that share at least one letter are not statistically distinguishable (*P* > 0.05), whereas groups with completely different letters are statistically distinguishable (*P* < 0.05). (**A**) The number of eggs laid per female is reduced when the mdn-LO neurons are selectively silenced by expression of the hyperpolarizing channel *Kir2.1* in the *tmc-GAL4* line. One-way ANOVA with Tukey’s Multiple Range Test. In this and the other figures: Groups that share at least one letter are not statistically distinguishable (*P* > 0.05), whereas groups with completely different letters are statistically distinguishable (*P* < 0.05). (**B)** The number of eggs laid per female of mdn-LO neurons selectively silenced, using Kir2.1, in the presence and absence of suppression by *elav-GAL80*. The number of eggs laid is restored by neuron-specific *elav-GAL80* suppression. One-way ANOVA with Tukey’s Multiple Range Test. (**C**) The egg-jamming ratio of *UAS-Kir2.1/+, tmc-GAL4/+*, *tmc-GAL4/+*;*UAS-Kir2.1/+,* and *tmc-GAL4/elav-GAL80*;*UAS-Kir2.1/+* flies. Fisher’s exact test. (**D**) Representative images of ovaries of *UAS-Kir2.1/+, tmc-GAL4/+*, *tmc-GAL4/+*;*UAS-Kir2.1/+,* and *tmc-GAL4/elav-GAL80*;*UAS-Kir2.1/+* flies. (**E**) Using the nuclear-localized RFP (UAS-Redstinger) and the membrane-localized GFP (UAS-mCD8::GFP; UAS-mCD8::GFP) as a marker, we investigated the labeling of mdn-LO neurons in the lateral oviducts in the presence of vGlut-GAL80 and elav-GAL80 strains. Both vGlut-GAL80 and elav-GAL80 can effectively block the tmc-GAL4 labeling of lateral oviduct neurons (mdn-LO) but not neurons in the CNS. Arrowheads indicate the mdn-LO neurons in the lateral oviducts. (**F**) Representative images of ovaries from *UAS-dTrpA1/+, tmc-GAL4/UAS-dTrpA1, elav-GAL80;tmc-GAL4/ UAS-dTrpA1*, and *tmc-GAL4,vGlut-GAL80/+; UAS-dTrpA1/+* flies at 22°C and 30°C. (**G**) The egg-jamming ratio of *UAS-dTrpA1/+, tmc-GAL4/UAS-dTrpA1*, *elav-GAL80;tmc-GAL4/ UAS-dTrpA1*, and *tmc-GAL4,vGlut-GAL80/+*; *UAS-dTrpA1/+* flies. Fisher’s exact test. (**H**) Representative images of *UAS-CsChrimsonmVenus*/+ and *tmc-GAL4/+*;*UAS-CsChrimsonmVenus/+* ovaries after optogenetic activation. The arrow indicates the location where the egg is jamming in the oviduct. (**I**) Optogenetic activation of *tmc-GAL4* neurons using CsChrimson. The egg-jamming ratio after optogenetic activation of *tmc-GAL4* neurons. Fisher’s exact test.

**Figure 3—figure supplement 1.**
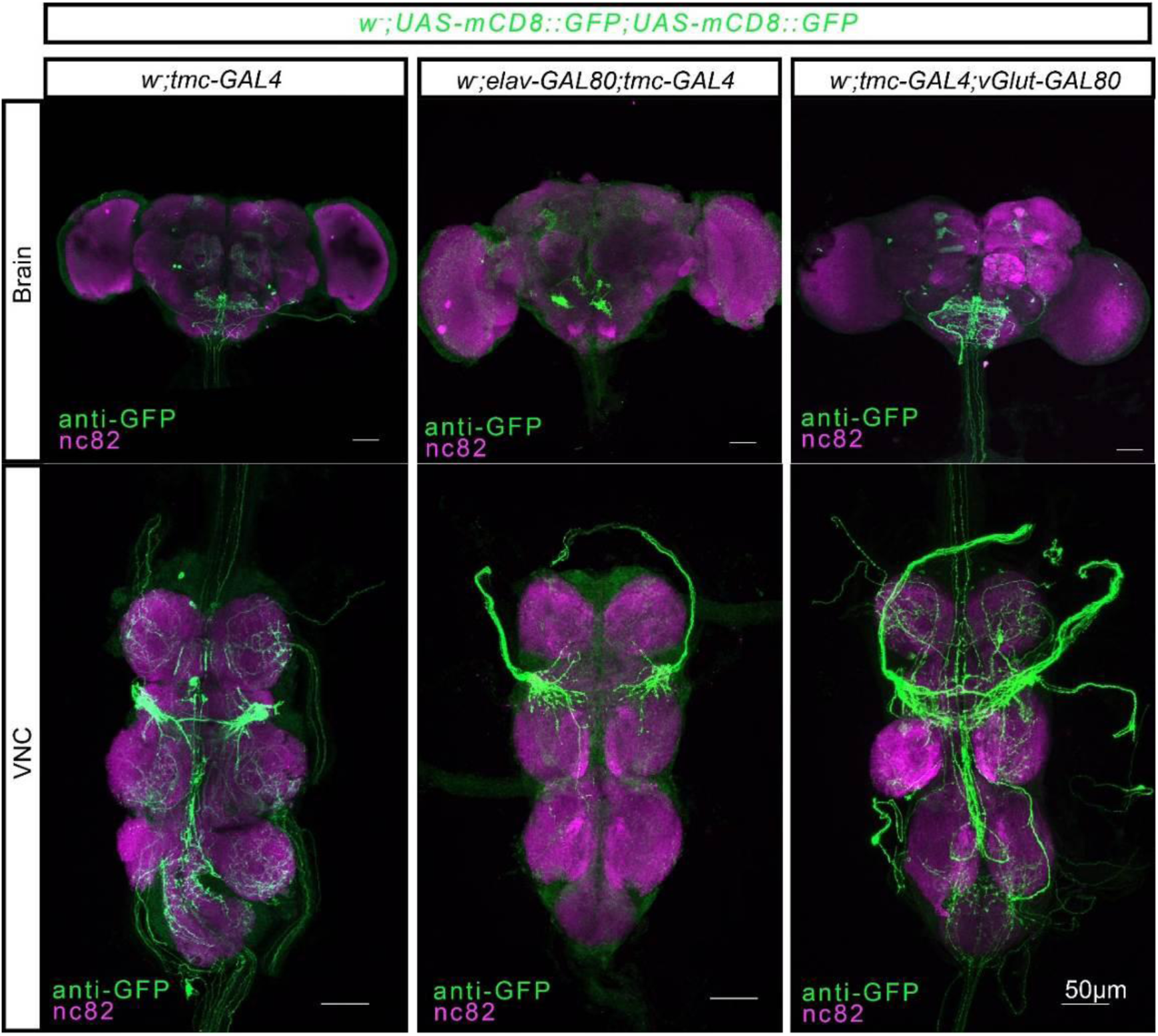
Expression of tmc-GAL4 in the presence of different GAL80 in the brain and ventral nerve cord. Using the membrane-localized GFP (*UAS-mCD8::GFP; UAS-mCD8::GFP*) as a marker, we investigated the labeling of mdn-LO neurons in the CNS in the presence of *vGlut-GAL80* and *elav-GAL80* strains. Note that in the *tmc-GAL4, elav-GAL80 > GFP*, only a subset of the GFP-labeled inputs to the subesophageal zone and the metathoracic ganglion of the VNC are retained; in the *tmc-GAL4; vGlut-GAL80 > GFP*, by contrast, there are fewer labeled axons in the ANm, indicating that inputs from the reproductive system are blocked.

Next, we selectively activated the TMC-expressing neurons thermo-genetically by using the *tmc-GAL4* to direct the expression of dTrpA1 <u>41</u>. Activation of *tmc*-GAL4 neurons via dTrpA1 (at 30°C) severely impaired oviposition and induced a significant egg-jamming rate (Figure 3F and G). To clarify which of the *tmc-GAL4* neurons are functionally required for coordinated oviposition and avoidance of egg-jamming, we suppressed the activity of GAL4 in selected tissues and cell types using the GAL80 repressor <u>40</u>. For this, we used two GAL80 lines, *elav-GAL80* <u>42</u> and vesicular glutamate transporter (*vGlut*)*-GAL80* <u>43</u>. We found that *elav-GAL80* and *vGlut-GAL80* lines both block the expression of TMC in mdn-LO neurons in the lateral oviduct (Figure 3E, Figure 3—figure supplement 1). Given that the motor neurons controlling muscle movements in *Drosophila* are glutamatergic <u>44</u>, *vGlut-GAL80* serves to repress any potential GAL4 miss-expression in motor neurons.

To determine whether the behavioral abnormality was due to the mdn-LO neurons, we combined the *elav*-*GAL80* or *vGlut*-*Gal80* transgene with the *tmc-GAL4* and *UAS-dTrpA1* to suppress activation of mdn-LO neurons. We found that animals with these manipulations showed no difference in egg-jamming compared with control flies, while *tmc-GAL4/UAS-dTrpA1* animals displayed increased egg-jamming at 30°C (Figure 3F and G).

To further confirm our findings, we used the optogenetic effector CsChrimson <u>45</u>, which affords greater control over the dynamic range of neuronal activation than *dTrpA1* <u>46</u>, and found that optogenetic stimulation of *tmc-GAL4* expressing neurons in *UAS-CsChrimsonVenus/tmc-GAL4* females abolished oviposition, dramatically different from all control females [see also <u>28</u>]. We also observed a significant egg-jamming rate in the activated *UAS-CsChrimsonVenus/tmc-GAL4* females (Figure 3H and I). Combining pan-neuronal expression of GAL80 with *UAS-CsChrimsonVenus/tmc-GAL4* significantly reduces the rate of egg-jamming seen after photoactivation (Figure 3I).

### *tmc*-expressing sensory neurons in the reproductive oviduct can sense contractions of the lateral oviduct

Our experiments, so far, suggest that the female flies sense the passage of eggs through the reproductive tract by means of mechanosensory neurons. Thus, we next asked whether the mdn-LO neurons monitor oviduct muscle contractions and oviposition.

To investigate whether ovulation and oviposition affect activity in mdn-LO neurons, we conducted an assay using a transcriptional reporter of intracellular Ca^2+^ (TRIC) that monitors sustained neuronal activity <u>47</u>. Indeed, we found a significant elevation of TRIC signals, corresponding to increased intracellular Ca^2+^ levels, in mdn-LO neurons of egg-laying virgin flies compared with virgin flies that did not lay eggs, monitored over 5 days (Figure 4A and A’). This finding is consistent with elevated long-term activity in mdn-LO neurons during egg laying.

**Figure 4.**
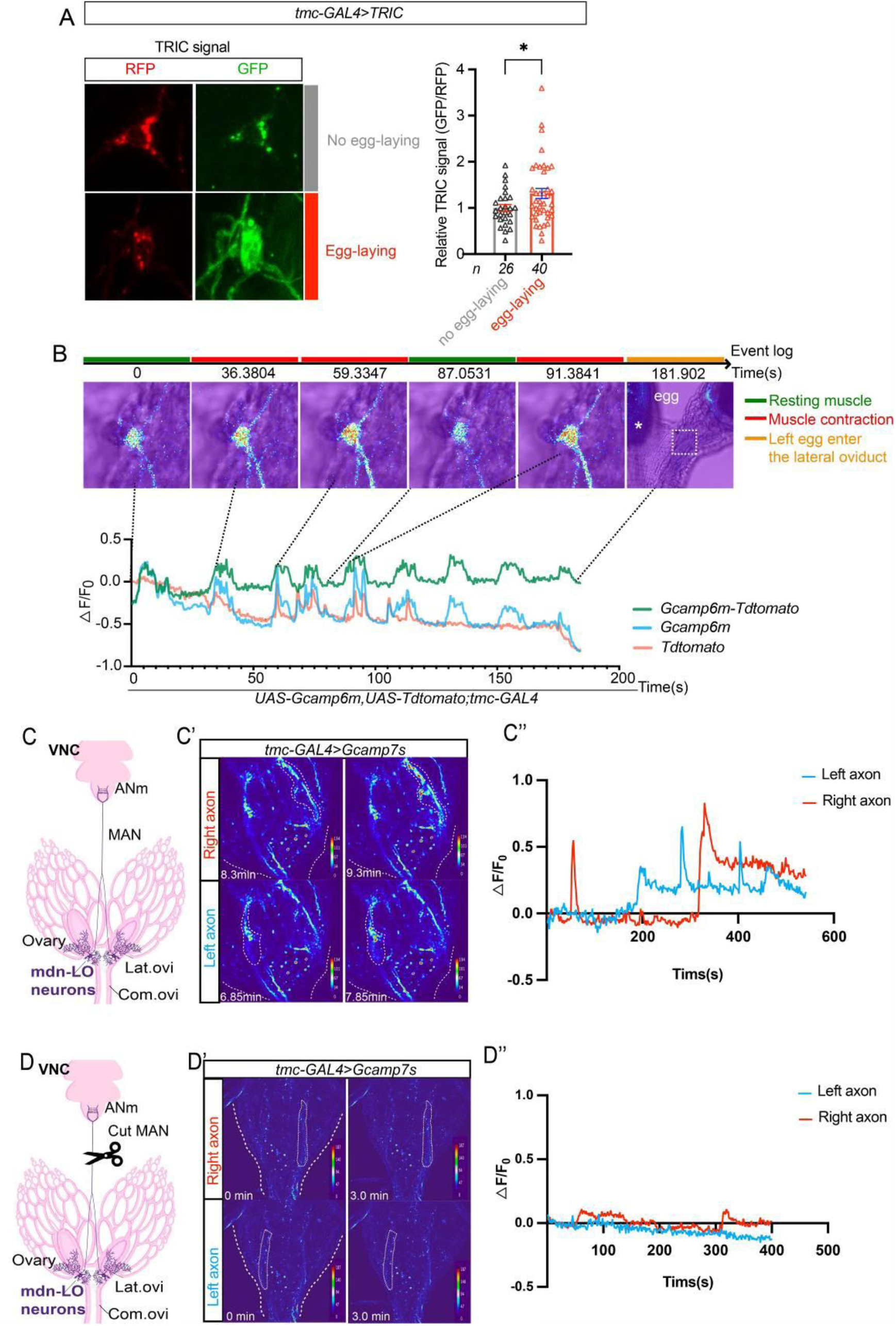
The lateral oviduct *tmc-*expressing neurons (mdn-LO) are mechanosensory and are activated during ovulation. (**A**) Monitoring sustained calcium activity in mdn-LO neurons using the TRIC method. Comparison of TRIC signals of mdn-LO sensory neurons between egg-laying and non-egg-laying females and representative images. We collected flies that were 5 days post-eclosion and dissected them, separating those that laid eggs and those that did not lay any eggs. After dissection, without staining, we immediately observed the TRIC signal of mdn-LO in real time under the same parameters (we used GFP/RFP as the TRIC signal). \**P* < 0.05, \*\**P* < 0.01, \*\*\**P* < 0.001, \*\*\*\**P* < 0.0001, and ns (non-significant); Two-tailed t-tests. (**B**) When the lateral oviduct contracts rhythmically, it is accompanied by rhythmic excitation of mdn-LO sensory neurons, as shown by increased Ca^2+^ levels monitored with Gcamp6m. (**C-C’’**) In intact animals (median abdominal nerve, MAN, intact), lateral oviduct mdn-LO sensory neurons transmit excitation to the abdominal neuromere (ANm), and Ca^2+^ activity in axon terminations of the left and right neurons can be monitored. (**D-D’’**) When the MAN is surgically severed, no Ca^2+^ activity is detected in axon terminations in the ANm during spontaneous muscle contractions. This supports that the activation in intact animals is derived from the axons of the lateral oviduct mdn-LO sensory neurons.

**Figure 4—figure supplement 1.**
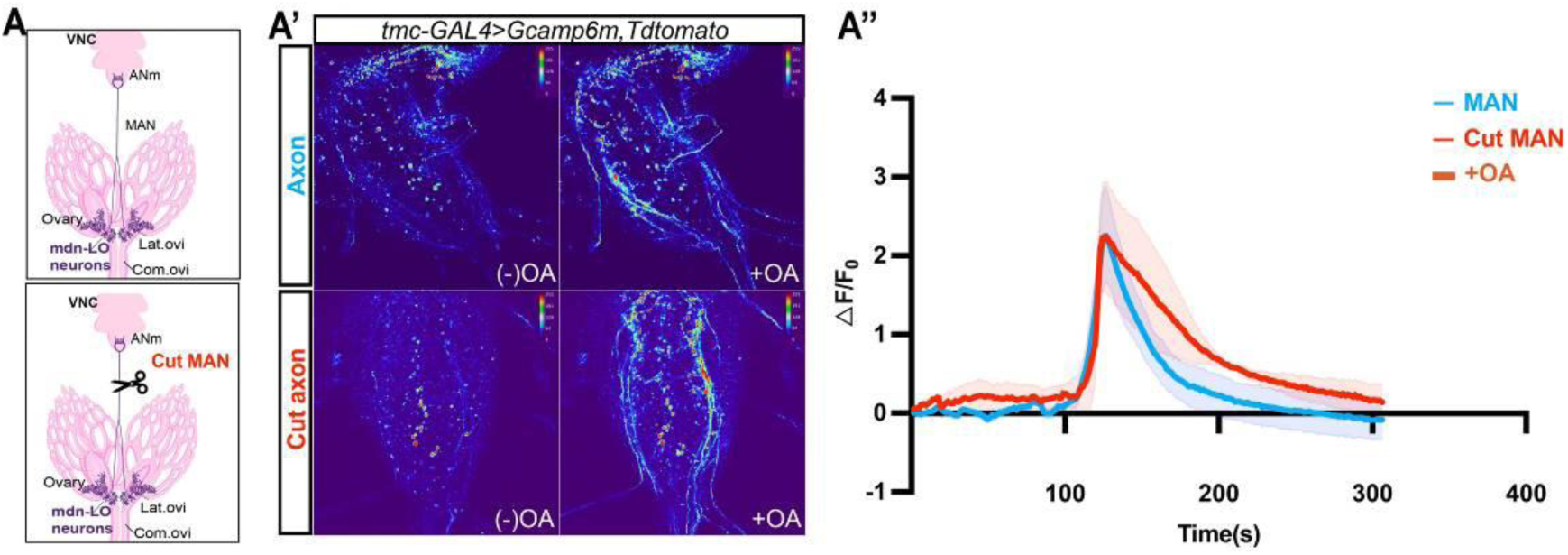
Mdn-LO neurons can be activated by octopamine in intact animals and after severing the connection to the oviduct. (A-A’’) *tmc*+ neurons can be activated by octopamine (A). Illustration of maintaining the connection via the median abdominal nerve (MAN) and cutting MAN. (A’) Calcium imaging of mdn-LO neuron axons in the ANm region before and after the addition of 1 mM octopamine in intact animals and after severing the MAN. (A’’) Changes in fluorescence intensity (△F/F_0_) of Gcamp6m in mdn-LO neuron axons in the ANm region before and after octopamine addition.

Next, to further explore the requirement of mdn-LO neurons for ovulation-induced neuronal activity, we monitored Ca^2+^ dynamics in the cell body of mdn-LO neurons in response to ovulation. To visualize changes in Ca^2+^ levels in cell bodies of mdn-LO, we expressed a transgene encoding the Ca^2+^ sensor GCaMP6m (*UAS-GCaMP6m*) and Tdtomato (*UAS-Tdtomato*) <u>48</u> in mdn-LO neurons under the control of the *tmc-GAL4.* We surgically exposed the lateral oviduct region of the female reproductive area, aiming to monitor the fluorescence changes of mdn-LO neurons as the egg passed through the lateral oviduct. However, we found that when the egg passes through the lateral oviduct, an inevitable movement occurs along the Z-axis, causing us to lose the signal. Even with the addition of tdTomato as a negative control, the tissue displacement still prevented us from accurately monitoring the change in fluorescence in the mdn-LO neurons as the egg passed through the lateral oviduct. Therefore, we monitored the activity of mdn-LO neurons when the lateral oviducts undergo spontaneous contractions. We observed that calcium activity in the cell body of the mdn-LO neurons increased when the muscle contracted (Figure 4B and Movie S1).

We also used another calcium sensor, GCaMP7s <u>49</u>, to monitor activity in axonal termini of mdn-LO neurons in the VNC. This was done since the GCaMP7s signal in *tmc-*expressing processes in the VNC is less likely to move out of the focal plane when oviduct contraction occurs, and thus facilitates measurements. We found that the CGaMP7s signal in the axonal termini of *tmc* neurons increased significantly during oviduct contraction (Figure 4C-C’’; Movie S2). To ensure that such GCaMP7 increase was not due to ‘‘local interaction’’ between *tmc-*expressing neurons and other neurons in the VNC, we physically severed the connection (median abdominal nerve, MAN) between the oviducts and the VNC. In these severed preparations, we did not observe a significant GCaMP7s increase in *tmc-*expressing axons in the VNC during oviduct contractions (Figure 4D-D’’; Movie S3). This finding supports that mdn-LO neurons convey contraction-related signals from the oviduct to the VNC rather than the Ca²⁺ transients being the result of local neuronal interactions in the VNC. Thus, our data strongly suggest that the *tmc*-expressing mdn-LO neurons of the lateral oviducts are mechanosensory and can detect contractions/distentions in this portion of the reproductive tract.

We next asked whether mdn-LO neurons can also be influenced by inputs other than mechanosensory stimulation during ovulation. Since octopamine (OA) has been implicated as a regulator of ovulation in insects <u>18</u>, we tested the effect of exogenous OA on mdn-LO activity. We found that OA application increased Ca²⁺ signals in the axon terminals of GCaMP6m-expressing mdn-LO neurons both in intact animals and after severing the connection to the oviduct (Figure 4 —figure supplement 1), suggesting OA modulates activity in mdn-LO neurons independently of mechanosensory input.

Finally, we used a chemical-genetic approach <u>50</u>, <u>51</u> to stimulate the mdn-LO neurons. We expressed the ATP-gated P2X2 channel, which is not expressed endogenously in flies, and can be used for ‘chemogenetic activation’ of specific neurons of interest. When stimulating the P2X2 expressing mdn-LO neurons with ATP, a robust contraction of the oviduct muscle was induced (Movie S4). This shows that mdn-LO mechanosensory neurons in the lateral oviduct can not only sense oviduct contraction, but also act through downstream motoneurons to regulate contractions of the lateral oviduct.

### ILP7-expressing neurons in the abdominal ganglion receive input from *tmc*-expressing mdn-LO neurons

We next sought to identify how TMC-expressing mechanosensory mdn-LO neurons engage the central circuitry that controls oviduct contractility.

To map candidate downstream partners of mdn-LO neurons, we first used trans-Tango <u>52</u> to visualize neurons postsynaptic to *tmc*-GAL4–labeled mdn-LO neurons. Expression of trans-Tango in mdn-LO neurons labeled numerous putative postsynaptic neurons, with the strongest labeling in the ABD of the VNC (Figure 5A-C’’) and in the suboesophageal zone (SEZ) (Figure 5—figure supplement 1A). We, however, did not detect a trans-Tango signal in any local neurons in the ovary or oviduct (Figure 5—figure supplement 1B). This finding further supports that mdn-LO neurons act as sensory neurons that transmit signals from the oviduct muscle to the central nervous system. Since the trans-Tango labeled numerous neurons, the identification of individual post-synaptic neurons was not possible. It has been shown previously <u>22</u>, and we confirmed here, that some of the *Ilp7-GAL4*-expressing neurons in the ventral portion of the posterior VNC are efferent neurons that innervate the oviduct (Figure 5D, Figure 5—figure supplement 2H). Importantly, it was previously shown that inhibiting ILP7*-*neurons consistently causes one or more eggs to be jammed in the oviduct <u>4</u>. Furthermore, efferent neurons in the posterior VNC that produce OA, identified by a *tdc2*-*GAL4,* are also involved in the regulation of ovulation <u>19</u>, <u>53</u>, <u>54</u>, <u>55</u>. Thus, we next used the syb:GRASP technique <u>56</u> to address whether axons of *tmc*-expressing mdn-LO neurons directly contact processes of *Ilp7*-expressing or *tdc2*-expressing neurons in the abdominal neuromeres of the VNC. In this method, half of the GFP is targeted specifically to the presynaptic termini of the mdn-LO neurons and the other to *Ilp7* or *Tdc* neurons, and GFP reconstitution occurs at exposure to the synaptic cleft only upon synaptic vesicle fusion. With *Ilp7* neurons and *Tdc* neurons separately as partners of mnd-LO neurons, we were able to detect reconstituted GFP clearly in the ABD of the VNC (Figure 5E-H; Figure 5—figure supplement 2B, D), suggesting that mdn-LO neurons can act as presynaptic partners of both *Ilp7* neurons and *Tdc* neurons in the ABD. However, we did not find evidence with the GRASP technique that *Ilp7* or *tdc2* neurons are coupled pre-synaptically to mdn-LO neurons (Figure 5—figure supplement 1C and 1E). Thus, the activation of mdn-LO neurons by applied OA, shown in Figure 5—figure supplement 1, is likely to represent paracrine (non-synaptic) signaling from octopaminergic neurons. Furthermore, we found that *tdc2-*GAL4 and *Ilp7*-GAL4 signals are not co-expressed in neurons in the ABD or the brain (Figure 5—figure supplement 1I). However, we could show by GRASP that *tdc2* neurons are presynaptically coupled to *Ilp7*-expressing neurons, suggesting functional connections between these neurons (Figure 5—figure supplement 1F-G).

**Figure 5.**
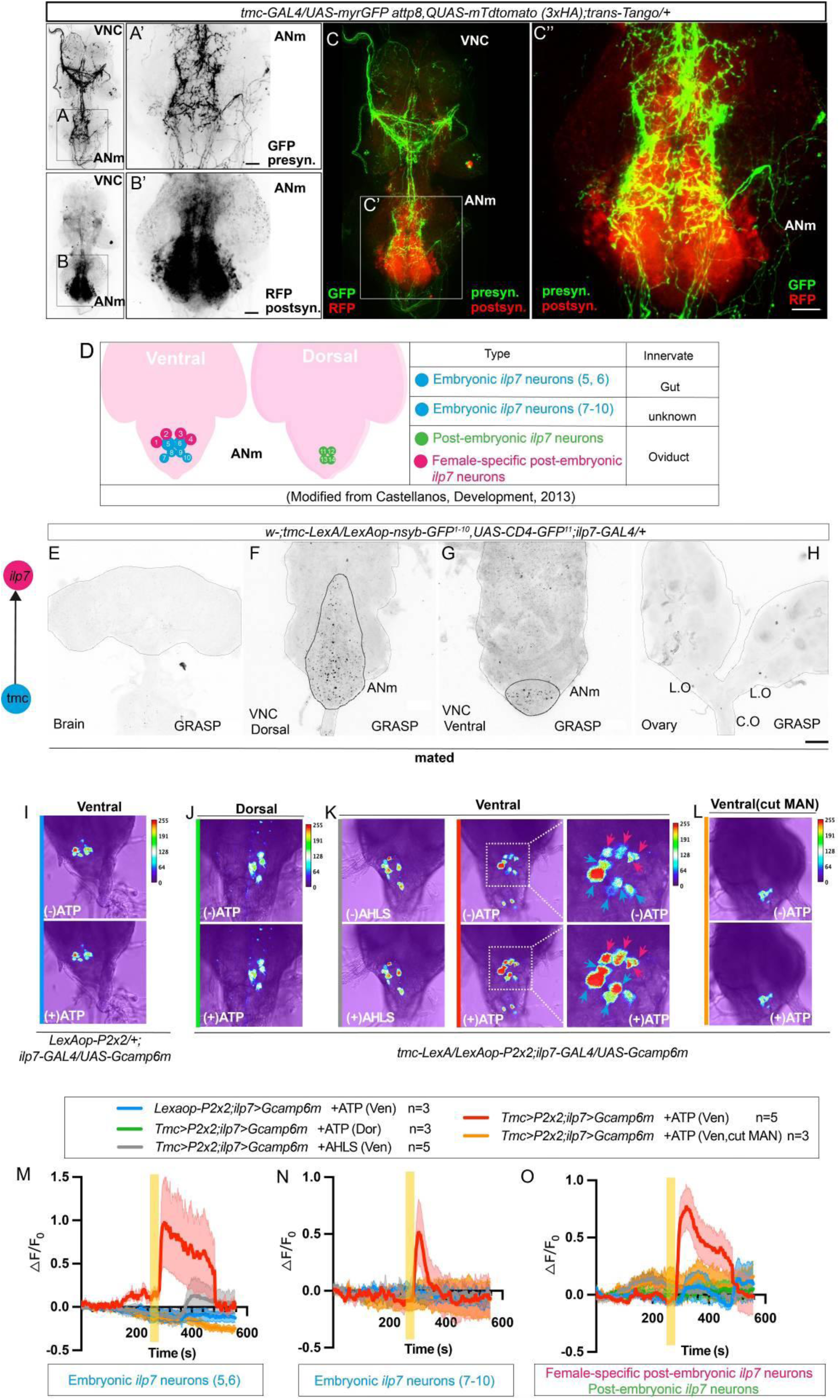
The mdn-LO sensory neurons regulate activity in the *Ilp7* neurons in the ANm, and their connections are visualized by trans-tango and GRASP. (A-C’’) Putative second-order neurons targeted by *tmc*+ neurons (green) in the ANm region revealed by trans-Tango (red) driven by *tmc*-GAL4. In A-B’, we show the expression of presynaptic signal (GFP/dark) in mdn-LO neurons and postsynaptic signal (RFP/dark) of *tmc*+ neurons in the ANm region of VNC. In C-C, merged confocal images of trans-Tango labeling of postsynaptic neurons (GFP/green) to *tmc*+ neurons (RFP/red). (D) Location of *Ilp7* neurons in the ventral and dorsal ANm regions. Blue circles indicate embryonic *Ilp7* neurons; pink circles indicate large female-specific post-embryonic *Ilp7* neurons; Green circles indicate the post-embryonic *Ilp7* neurons. Original data from <u>22</u>. The three types of *Ilp7 –*expressing cell clusters innervate different tissues. The embryonic *Ilp7* neurons innervate the intestine, and the female-specific post-embryonic *Ilp7* neurons and post-embryonic *Ilp7* neurons are glutamatergic motoneurons that innervate the oviduct muscle of the female reproductive system. (E-H) GRASP signals between mdn-LO sensory neurons and *Ilp7* neurons are observed in the brain, dorsal and ventral VNC, and ovary. (I–L) The mdn-LO sensory neurons regulate the excitation of ventral VNC Ilp7 neurons in the ANm region. In control preparations lacking P2X2 expression in mdn-LO neurons (LexAop-P2X2/+), application of 1 mM ATP did not increase Ca²⁺ activity in ventral Ilp7 neurons (I). In contrast, activating P2X2-expressing mdn-LO neurons with 1 mM ATP robustly increased Ca²⁺ activity in ventral Ilp7 neurons, whereas dorsal Ilp7 neurons were not affected (J). (K) Vehicle control: AHLS (ATP solvent) applied with the same timing/volume elicited no response. (L) MAN-severed preparation: after cutting the median abdominal nerve (MAN) connecting the reproductive tract/periphery to the abdominal ganglion, ATP application failed to evoke ventral Ilp7 responses. (M-O) Graphs showing activation of *Tmc*+ neurons via *P2X2* increased different *Ilp7+* motor neuron activity. Fluorescence changes (ΔF/F₀) of *GCaMP6s* in *Ilp7* neurons are shown following different treatments. The yellow bar indicates the addition of the solution.

**Figure 5—figure supplement 1.**
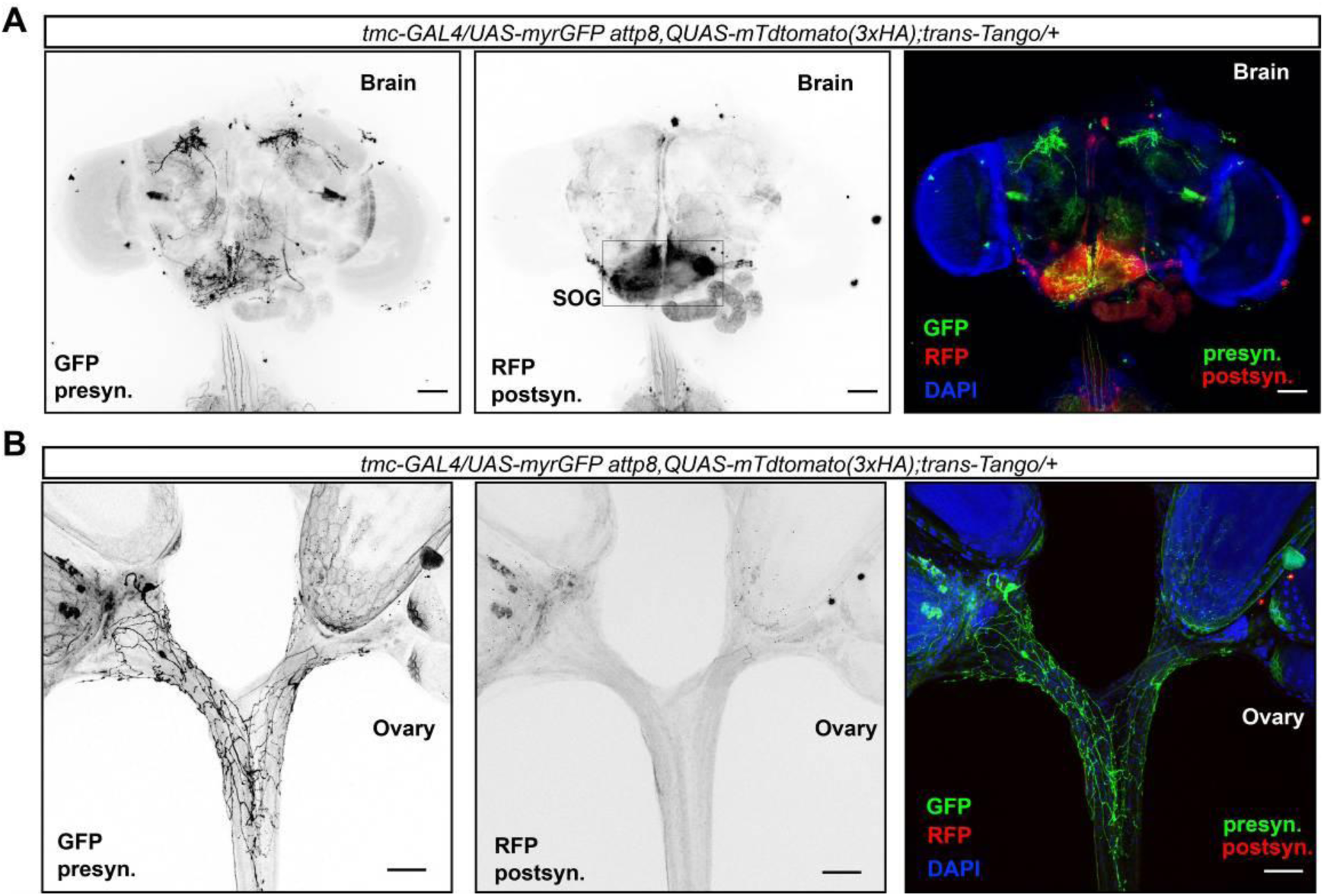
The presumed second-order neurons of mdn-LO target the subesophageal zone (SEZ) region in the brain. (A) Putative second-order neurons of *tmc*+ neurons (green) in the suboesophageal ganglion (SOG) region revealed by trans-Tango (red). (B) Using the trans-tango system, it was found that there are no putative second-order neurons of mdn-LO in the oviduct.

**Figure 5—figure supplement 2.**
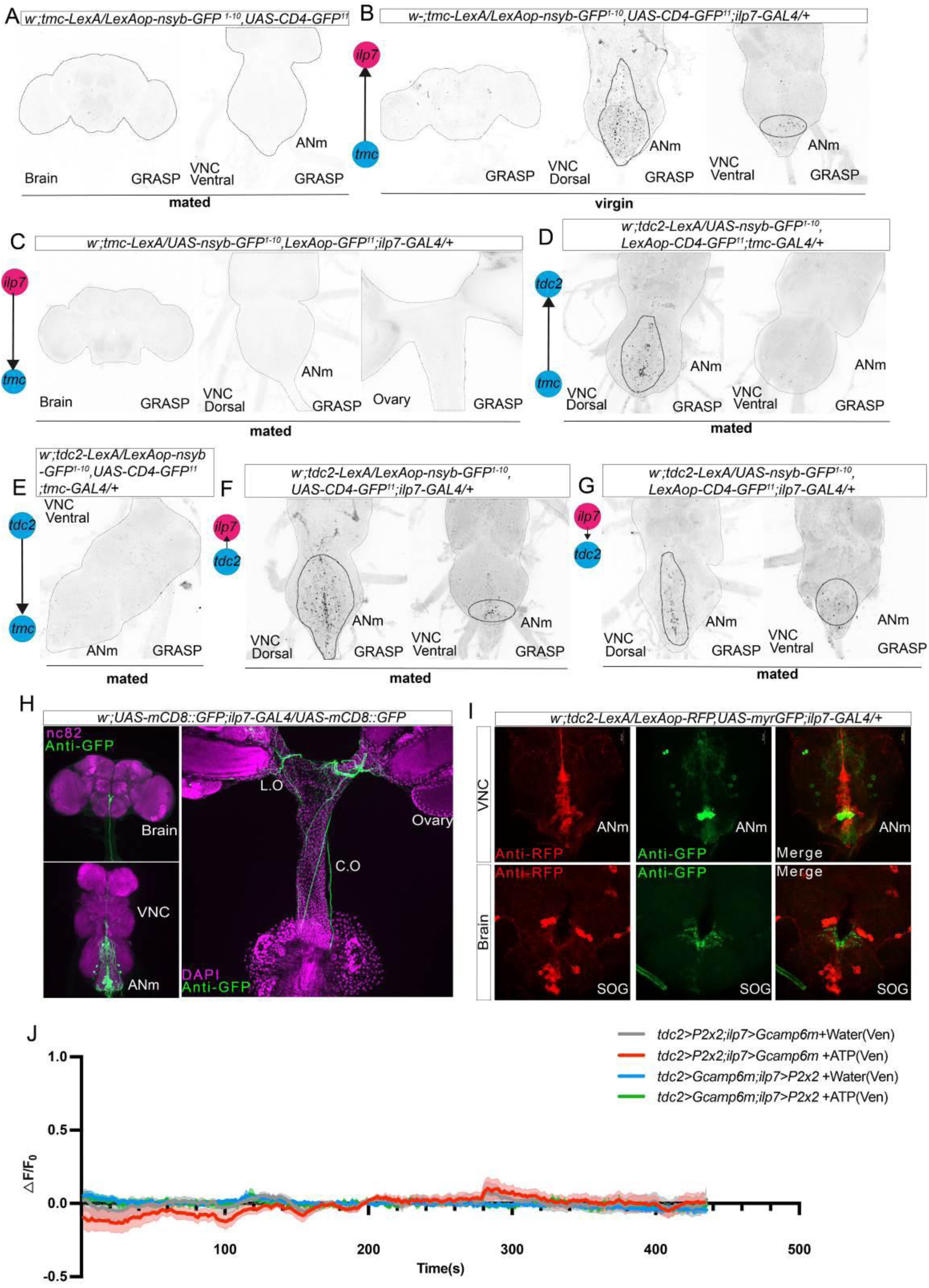
Probing connections between *tdc2*+, *Ilp7*+, and *tmc*+ neurons by the GRASP technique and calcium imaging. (A) Lack of GRASP signal from expressing *tmc-LexA* alone in the brain and ventral VNC. (B) Flies in virgin status were used to obtain the GRASP signal between mdn-LO (expressing *tmc-LexA*) and *Ilp7* + neurons with *Ilp7-GAL4* in the brain, dorsal, and ventral VNC. The black signal represents the region of putative synaptic connections (indicated by black dashed lines). (C) No GRASP signal is seen from the *Ilp7* + neurons to *tmc*+ neurons in the brain, dorsal VNC, and ovary. (D) GRASP signal from the *tmc*+ neurons to *tdc2*+ neurons in the dorsal and ventral VNC. The dark signal represents the region of synaptic connections (indicated by black dashed lines). (E) No GRASP signal is seen from the *tdc2*+ neurons to *tmc*+ neurons in the ventral VNC. (F) GRASP signal from the *tdc2*+ neurons to *Ilp7* + neurons in the ventral and dorsal VNC. The dark signal represents the region of putative synaptic connections (indicated by black dashed lines). (G) GRASP signal from the *Ilp7* + neurons to *tdc2*+ neurons in the ventral and dorsal VNC. The dark signal represents the region of putative synaptic connections (indicated by black dashed lines). (H) The expression of *Ilp7*-GAL4 positive neurons in the brain, VNC, and ovary. (I) *Ilp7*+ (green) and *tdc2*+ (red) neurons within the ANm and brain are distinct. (J) There is no functional connection between ILP7 neurons and Tdc2 neurons. When ILP7 neurons are activated by ATP, the calcium signal in Tdc2 neurons shows no significant change; the converse is also true.

To test whether the contacts observed between mdn-LO and ILP7 neurons using the GRASP technique permit functional signal transmission, we next asked whether stimulation of mdn-LO neurons affects Ca²⁺ levels in ILP7 neurons. For this, we established an *ex vivo* “brain-less” preparation in which the full oviduct, where ovaries remain connected to the VNC, while the connection to the brain is severed. In this preparation, we monitored Ca²⁺ dynamics in ILP7 neurons using GCaMP6m <u>48</u>. To stimulate mdn-LO neurons, we expressed in these neurons the ATP-gated P2X2 channel <u>50</u>, <u>51</u>.

We found that stimulating the P2X2-expressing mdn-LO neurons with ATP triggered a significant increase in GCaMP signal in the cell bodies of *Ilp7* neurons in the ventral portion of the abdominal neuromeres of the VNC (Figure 5I-O), accompanied by rhythmic contractions of the oviduct (Movie S4). We performed three control experiments where no increase in GCaMP signals was observed in ILP7 neurons: (i) genotype controls lacking P2X2 expression in mdn-LO neurons (Figure 5I), (ii) vehicle controls, AHLS (ATP solvent) applied with the same timing/volume elicited no response (Figure 5K), and (iii) preparations in which the median abdominal nerve that connects oviduct and VNC was severed (Figure 5L).

Interestingly, we did not detect comparable elevations of the GCaMP signal in ILP7 neurons located on the dorsal side of the ABD. Thus, the Ca²⁺ increase was confined to the ventral cluster that includes the previously identified female-specific post-embryonic ILP7 oviduct motoneurons (Figure 5D, 5K, 5M-5O). Collectively, our findings demonstrate that mdn-LO axons form functional contacts with ILP7 neurons in the ABD, and that activation of the mdn-LO neurons induces increased Ca²⁺ activity in the ventral subset of these ILP7 neurons, which in turn modulates contractions in the lateral oviducts.

### ILP7 neuron activity disrupts egg passage and exhibits mechanically tunable rhythmic Ca²⁺ dynamics

Previous work showed that hyperpolarizing *Ilp7-GAL4* neurons in the posterior VNC leads to egg jamming in the oviduct <u>57</u>. However, the effects of activating these neurons and the role of ILP7 peptide on ovulation and oviposition have not been investigated. We found that optogenetic activation of *Ilp7-GAL4* neurons using CsChrimson markedly increased the fraction of females exhibiting egg-jamming compared with genetic controls (Figure 6A). Furthermore, thermogenetic activation of *Ilp7-GAL4* neurons by driving dTrpA1 reduced the number of eggs laid at 30 °C compared to 22 °C controls and increased the incidence of egg jamming (Figure 6B–D).

**Figure 6.**
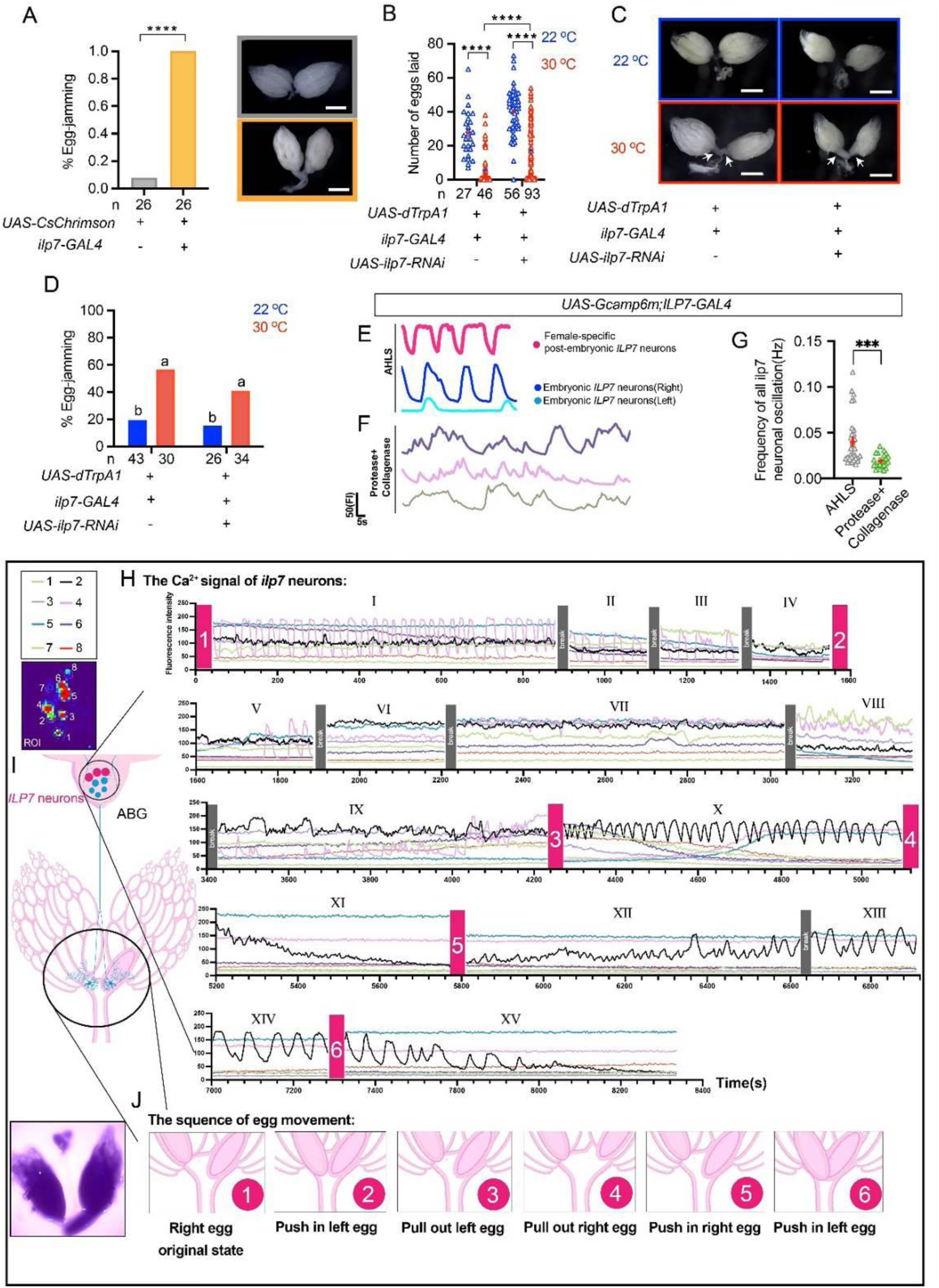
ILP7 neuron activity disrupts egg passage and exhibits mechanically tunable rhythmic Ca²⁺ dynamics. (A) Optogenetic activation of ILP7-GAL4 neurons (UAS-CsChrimsonmVenus; ILP7-GAL4) increases the fraction of females exhibiting egg jamming (left). Representative images of dissected female reproductive tracts after optogenetic stimulation are shown (right); the boxed example highlights a jammed egg in the oviduct. (B) Thermogenetic activation of ILP7 neurons (UAS-dTrpA1; ILP7-GAL4) reduces oviposition at 30 °C relative to 22 °C controls, and this reduction is partially alleviated by Ilp7 RNAi during activation (UAS-Ilp7-RNAi; UAS-dTrpA1; ILP7-GAL4). (C) Representative images of dissected female reproductive tracts from the thermogenetic activation experiment in (B) at 22 °C and 30 °C, showing egg retention/jamming in activated conditions (arrowheads). (D) Egg-jamming frequency following thermogenetic activation of ILP7 neurons, with or without Ilp7 RNAi, at 22 °C and 30 °C (genotypes as in B). Groups sharing at least one letter are not statistically distinguishable. (E) Spontaneous rhythmic Ca²⁺ oscillations recorded with GCaMP6m in ILP7-GAL4 neurons in the adult VNC preparation, including female-specific post-embryonic ILP7 neurons (pink) and embryonic ILP7 neurons (dark and light blue). (F) Rhythmic Ca²⁺ oscillations persist in ILP7 neurons after enzymatic digestion (protease/collagenase), although oscillations are slowed. (G) Quantification of oscillation frequency in ILP7 neurons before and after enzymatic digestion (AHLS, Adult Hemolymph-Like Saline). (H) ILP7-neuron Ca²⁺ dynamics are modulated by egg position within the oviduct. Traces show Ca²⁺ signals from eight ILP7 neurons (color-coded ROIs) across a continuous recording that was segmented into 15 consecutive time windows (roman numerals I–XV). Red numbered boxes indicate periods corresponding to discrete egg-position states defined in (J), whereas gray boxes mark recording intervals during which no egg manipulation was performed. In the initial state G1 (I–IV; right egg enters the oviduct), neuron 4 exhibits rhythmic Ca²⁺ oscillations that progressively slow and terminate by IV. After the left egg is pushed into the oviduct (G2, V–IX), neuron 4—previously quiescent—shows a brief burst of oscillations, then pauses again, with oscillations re-emerging by IX. After the left egg is pulled out (G3, X), neurons 1, 6, and 8 decrease in baseline signal, neuron 4 stops oscillating, and neurons 4 and 5 show increased activity; notably, neuron 2 displays a newly emergent oscillatory pattern. When the right egg is pulled out (G4, XI), neuron 2 activity decreases, whereas pushing the right egg back in (G5, XII–XIV) restores and re-initiates rhythmic oscillations in neuron 2, consistent with regulation by right-side mechanosensory feedback. Finally, when the left egg is pushed back into the oviduct (G6, XV), oscillations in neuron 2 disappear. (I) Schematic of the ex vivo imaging configuration used for the recording in (H), in which the ANm region of the VNC was retained for Ca²⁺ imaging. The upper inset indicates the eight ILP7 neurons monitored and the corresponding ROI color codes used in (H). (J) Schematic timeline of egg-position manipulations and scored egg-location states. Numbers indicate the sequence of events and correspond to the red numbered boxes in (H).

**Figure 7.**
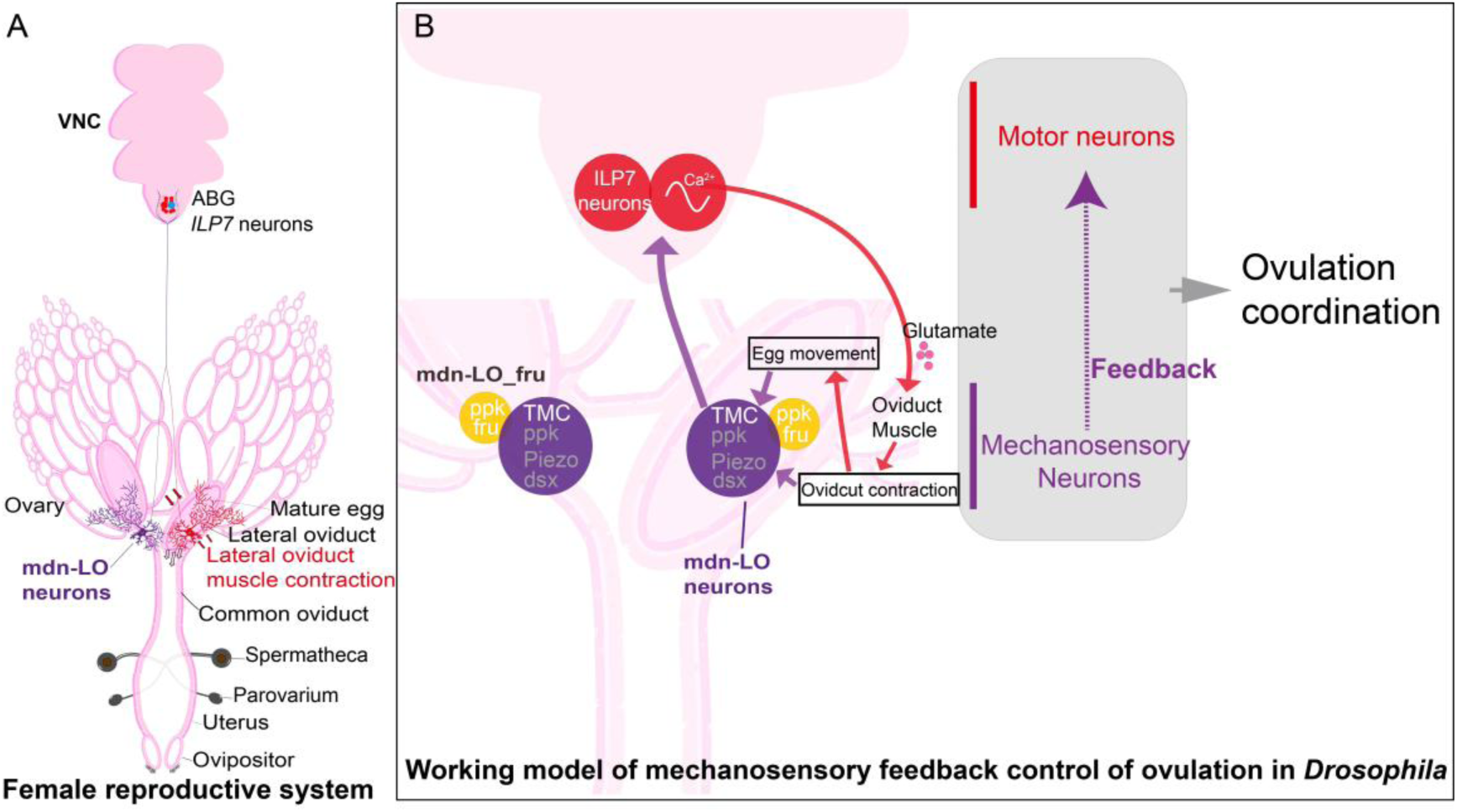
Proposed model of mechanosensory feedback control of ovulation in Drosophila. (A) Graphic summary of the female reproductive tract and ventral nerve cord (VNC). (B) We propose a working model of ovulation in which TMC-dependent lateral-oviduct mechanosensory neurons (mdn-LO; TMC/PPK/Piezo⁺) detect contraction– and egg-movement–linked mechanical cues and relay this information to the abdominal ganglion, where it shapes motoneuron activity patterns of ILP7-expressing glutamatergic oviduct motoneurons to coordinate egg passage from the paired lateral oviducts into the common oviduct and thereby promote successful ovulation. ILP7 peptide signaling contributes to overall oviposition output but is not a determinant of egg-jamming phenotypes under our experimental conditions. The yellow ppk⁺/fru⁺ LO neuron (here labeled mdn-LO_fru for clarity; previously described without a specific name). The mdn-LO_fru neurons constitute a second pair of lateral-oviduct multidendritic neurons (ppk⁺ fru⁺) reported previously; their role in egg-traffic coordination was not directly examined here.

To assess the contribution of the ILP7 neuropeptide during ILP7 neuron activation, we reduced ILP7 levels by RNAi (*Ilp7-GAL4 > UAS-Ilp7-RNAi*) while activating ILP7 neurons with dTrpA1. Under these conditions, the number of eggs laid was partially restored relative to ILP7 activation alone (Figure 6B), whereas the egg-jamming frequency was not significantly reduced (Figure 6D). Together, these data indicate that elevating neuronal activity in ILP7 neurons elicits egg-jamming, while the ILP7 peptide contributes primarily to overall oviposition rather than playing a role in the jamming phenotype (at least under our assay conditions). This finding might suggest that the ILP7 peptide acts at the level of the common oviduct rather than on the lateral ones.

We next examined the activity dynamics of ILP7 neurons using *Ilp7-GAL4 > GCaMP6m*. In *ex vivo* “brain-free” VNC preparations, *Ilp7*-GAL4–expressing female-specific post-embryonic neurons and embryonic ILP7 neurons display spontaneous rhythmic Ca²⁺ oscillations (Figure 6E). These two neuron types can be distinguished by their locations.

Following enzymatic digestion (protease/collagenase), the rhythmic oscillations were still observed, although they were slower (Figure 6F). Because digestion makes it difficult to classify ILP7 subtypes by position, we pooled the ILP7 neurons for quantification. The oscillation frequency decreased from ∼0.040 Hz before digestion to ∼0.019 Hz after digestion (Figure 6G), consistent with an intrinsic or locally generated oscillatory capacity, although additional network contributions to the rhythmicity cannot be excluded.

Finally, we asked whether the rhythmic activity of the ILP7-nerons is modulated by oviduct-linked internal mechanosensory feedback. Thus, we recorded prolonged Ca²⁺ activity from multiple ILP7 neurons while scoring egg position and movement within the reproductive tract. Across such a long recording, ILP7 neurons showed distinct oscillatory modes and transitions that covaried with changes in egg movement (Figure 6H). Time points at which egg position was scored are indicated (Figure 6H), and the corresponding sequence of egg movement states is summarized schematically (Figure 6I, J). Our observations suggest an association between the movement of the egg through the oviduct and Ca²⁺ dynamics in ILP7 neurons, and furthermore provide an indication of how internal mechanosensory feedback influences excitatory motor output during egg transport.

## Discussion

In this study, we describe a novel neuronal circuit that regulates coordinated oviposition in *Drosophila*. A pair of multidendritic sensory neurons, mnd-LO, in the lateral oviducts respond to muscle contractions as the egg passes and provide feedback to bilateral sets of motoneurons (ILP7 neurons) in the VNC to ensure coordinating lateral oviduct contractions to propel eggs into the common oviduct without jamming. In this circuit, we discovered a previously unrecognized role for the mechanosensory channel protein TMC. The TMC channel is expressed in the mnd-LO neurons together with two additional mechanosensory channels, PPK and PIEZO. Mutations in all three channel genes result in defects in ovulation/oviposition. However, only mutation or cell-specific knockdown of *tmc* markedly elevates the incidence of egg jamming at the junction between the two lateral oviducts and the common oviduct. Together, these findings highlight a key role for mechanosensory feedback in ensuring reliable egg transport in the reproductive tract, with TMC being specifically required to prevent jamming at the entrance to the common oviduct. Thus, while mdn-LO neurons co-express TMC, PPK, and PIEZO, our data suggest that these channels contribute in distinct ways to the mechanosensation of the lateral oviduct and that they function complementarily in this sensory process. Future work is required to understand in detail the specific roles of Piezo and PPK in the *Tmc*-positive sensory neurons. Interestingly, a recent report showed that PIEZO channels are required for effective uterine contractions during mammalian parturition <u>58</u>, suggesting an evolutionary conservation of function.

The mdn-LO neurons display rhythmic [Ca²⁺] activity during spontaneous and induced contractions of the oviduct muscle, consistent with a mechanosensory role. Using trans-Tango and GRASP, we found that mdn-LO neurons are presynaptic to a set of female-specific efferent neurons in the abdominal neuromere that co-express glutamate and the neuropeptide ILP7. Chemogenetic stimulation of mdn-LO neurons increases [Ca²⁺] in ILP7 neurons and elicits rhythmic contractions of oviduct musculature. Previous studies established that activity in these female-specific ILP7 neurons is required for proper ovulation and oviposition. It is likely mediated via glutamatergic neuromuscular transmission <u>4</u>, <u>8</u>, <u>19</u>, <u>22</u>. However, the functional contribution of the co-localized ILP7 peptide to oviduct contractions was not investigated. Our study demonstrates that blocking ILP7 peptide release (by *Ilp7*-RNAi) during activation of ILP7 neurons recovers oviposition but does not affect egg jamming. Thus, ILP7 may play a modulatory role further down the reproductive tract, such as in the common oviduct, and thereby influence the rate of egg laying, possibly by means of short-range hormonal signaling (or paracrine/non-synaptic signaling). In contrast, it is likely that the ILP7 neurons utilize glutamate for direct motor output to coordinate lateral oviduct muscle<u>4</u>, <u>8</u>, <u>19</u>, <u>22</u>.

Interestingly, the specific location of egg jamming in the oviduct depends on which component of the pathway is manipulated. Activation or inactivation of ILP7 neurons leads to egg jamming in the entrance of the common oviduct, whereas activation of mdn-LO neurons preferentially generates egg jamming in the lateral oviducts. This finding suggests that the two mdn-LO neurons provide sensory feedback from each of the lateral oviducts to modulate the two pairs of ILP7*-*motoneurons to generate alternating activity in the two lateral oviducts to ensure a smooth passage of eggs into the common oviduct.

Apart from the ILP7-neurons and mdn-LO neurons shown here and descending neuronal pathways from the brain <u>3</u>, <u>16</u>, there are additional neurons in the control of ovulation in *Drosophila*. These are a second pair of multidendritic sensory neurons in the lateral oviducts that express PPK, but not the TMC and PIEZO channels <u>8</u>, and a small set of octopaminergic efferent neurons (*tdc*-neurons) in the abdominal neuromeres of the VNC that innervate the muscle of the oviduct <u>17</u>, <u>18</u>, <u>19</u>. We did not specifically address the function of this second set of sensory neurons, but did investigate the possible interactions between mdn-LOs, ILP7*-*neurons, and the *tdc2*-expressing octopaminergic neurons. Using GRASP technique to reveal synaptic partners, we could not demonstrate inputs from mdn-LO neurons to *tdc*-expressing neurons or vice versa. However, we found that OA applied to the abdominal neuromere induced [Ca^2+^] increases in the axon terminations of the mdn-LO neurons. Since GRASP analysis could not visualize any synaptic contacts from *tdc2* neurons onto mdn-LOs, we suggest that OA acts after paracrine release (non-synaptic action) to modulate these sensory cells. By severing the axons of the *tdc2* neurons by cutting the nerve from the VNC to the oviduct, we could demonstrate that this OA action on mdn-LOs occurs locally in the VNC. Furthermore, using GRASP, we found that the *tdc2*-neurons are presynaptic to the ILP7 neurons. Hence, the *tdc2*-neurons, which are known triggers of ovulation <u>17</u>, <u>18</u>, <u>19</u>, may also trigger activity in the ILP7-neurons. In summary, we found no evidence that ILP7– and *tdc2-*neurons signal synaptically to mdn-LO neurons, but *tdc2-*neurons may synapse onto ILP7-neurons and also modulate mdn-LOs by paracrine signaling. Thus, while OA released from *tdc2-*neurons is a main trigger of ovulation, <u>17</u>, <u>18</u>, <u>19</u>, it may also be a signal to activate ILP7-neurons and to modulate the activity of mdn-LOs. Interestingly, the mdn-LOs do not provide mechanosensory feedback to *tdc2*-neurons, only the ILP7 neurons. Finally, the mechanistic role of the octopaminergic modulation of the mdn-LOs was not revealed in our study.

While our study delineates key elements of the regulatory ovulation pathway, the complete neuronal connectivity and state-dependent recruitment of additional premotor and interneuron partners remain to be described. For the octopaminergic neurons in the VNC that initiate ovulation, a recent screen of membrane receptors has identified regulatory inputs including ILP8, GABA, and SIFamide <u>59</u>. Similarly, single-cell transcriptomic datasets could be leveraged to search for candidate receptors and ion channels expressed by ILP7 neurons and mdn-LO neurons, thereby generating testable hypotheses for future functional validation. The emergence of comprehensive female adult VNC connectomic resources will further enable circuit-level tests of this working model by identifying additional mechanosensory inputs, premotor partners, and connectivity motifs associated with ILP7 neurons.

Notably, ILP7 neurons exhibit spontaneous rhythmic [Ca²⁺] dynamics in *ex vivo* preparations, and these dynamics are influenced by mechanical feedback linked to egg movement. Establishing how this rhythmicity relates to contractions in the oviduct muscle *in vivo* will benefit from future experiments that directly assay phase relationships and causal coupling under intact physiological conditions.

The precise role of ILP7 in the ovulation pathway remains to be clarified since we only found a possible role in sustaining oviposition. Previous studies detected no effect on ovulation of *Ilp7*-knockdown in the ILP7*-*neurons <u>4</u>, <u>22</u>. However, overexpression of *Ilp7* in ILP7-neurons, or ubiquitously, led to flies that were more susceptible to laying eggs on a high sucrose medium than a regular medium <u>4</u>. It is not clear how ILP7 mediates this effect, but it should be noted that apart from the sex-specific ILP7-neurons, there are further ILP7-expressing neurons (in both sexes) in the VNC of flies that may play roles in oviposition-related behavior and its relation to gut function and nutritional state <u>22</u>, <u>60</u>. Two of these are ascending interneurons that co-express short neuropeptide F (sNPF) and supply axons to the brain <u>60</u>, <u>61</u>, where they act on the insulin-producing cells (IPCs) to regulate insulin-signaling when flies ingest high-carbohydrate medium <u>62</u>, <u>63</u>. While we only investigated the role of the efferent oviduct-innervating ILP7-neurons (with ILP7 as a putative co-transmitter of glutamate), it may be interesting for the future to determine whether these and the ascending ILP7/sNPF neurons are functionally coupled and thereby link nutrition and oviposition.

Finally, the contribution of ovulation-related sensory input to the ramping activity of OviDN neurons that initiate egg deposition <u>3</u> warrants further investigation. A possible hypothesis is that the initial mechanosensory signals generated as an egg enters the oviduct or uterus are relayed, via as-yet-unknown pathways, to the brain, thereby prompting the fly to initiate the oviposition evaluation program and triggering the onset of calcium ramping in OviDNs. During this ramping phase, OviDNs may integrate external environmental cues and internal state signals until activity reaches a threshold, at which point the oviposition motor sequence is engaged to complete egg deposition. Therefore, elucidating the primary sensory circuits that trigger the oviposition evaluation program may enable the development of a model linking interoceptive sensory inputs to behavioral decision-making.

## Materials and methods

### Animals

Unless otherwise mentioned, all fly stocks were raised in standard cornmeal–molasses–agar medium maintained at 25°C, 60% humidity, and a 12 hr:12 hr light: dark cycle. We used *w^1118^* flies as the wild-type strain.

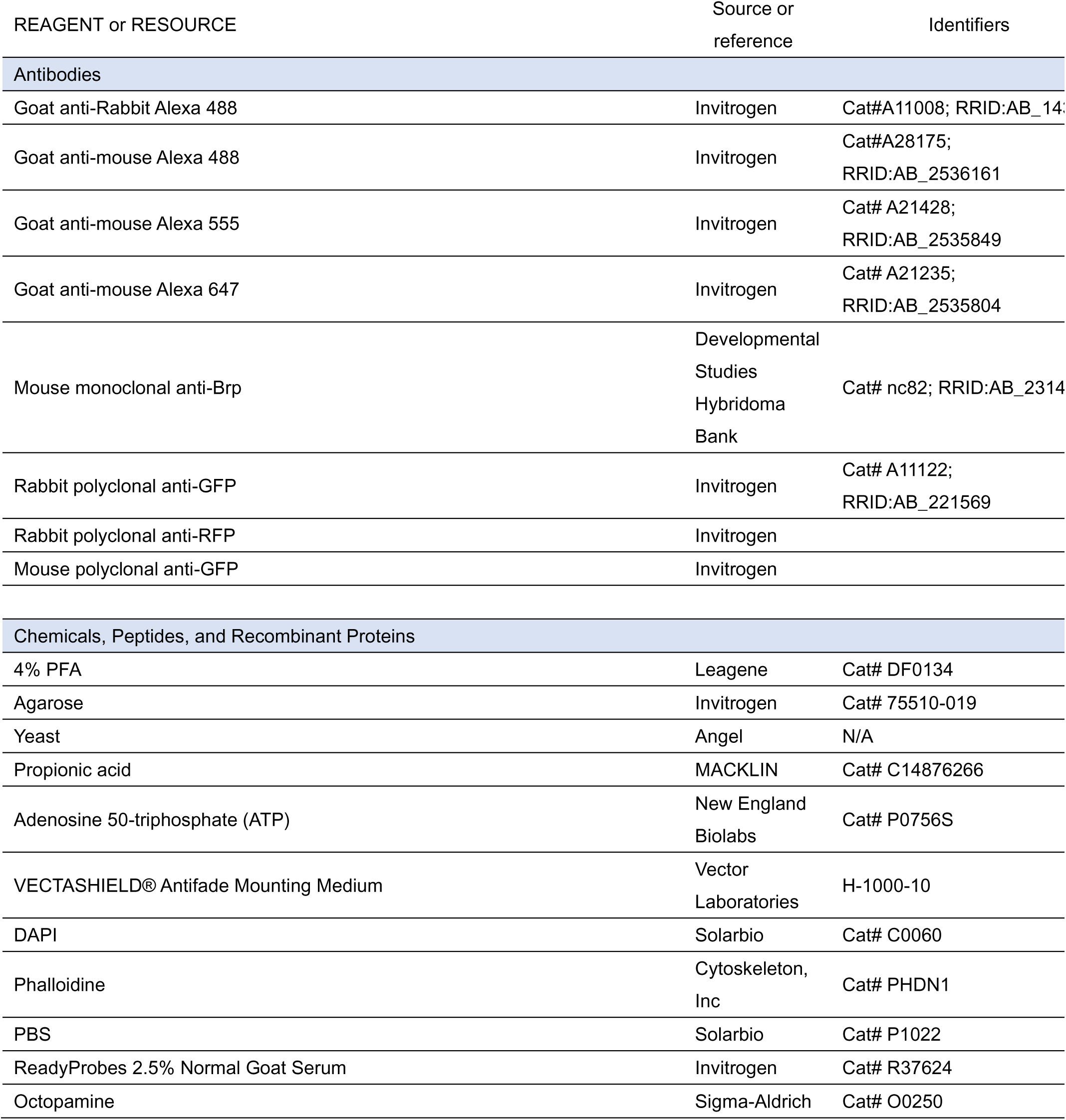
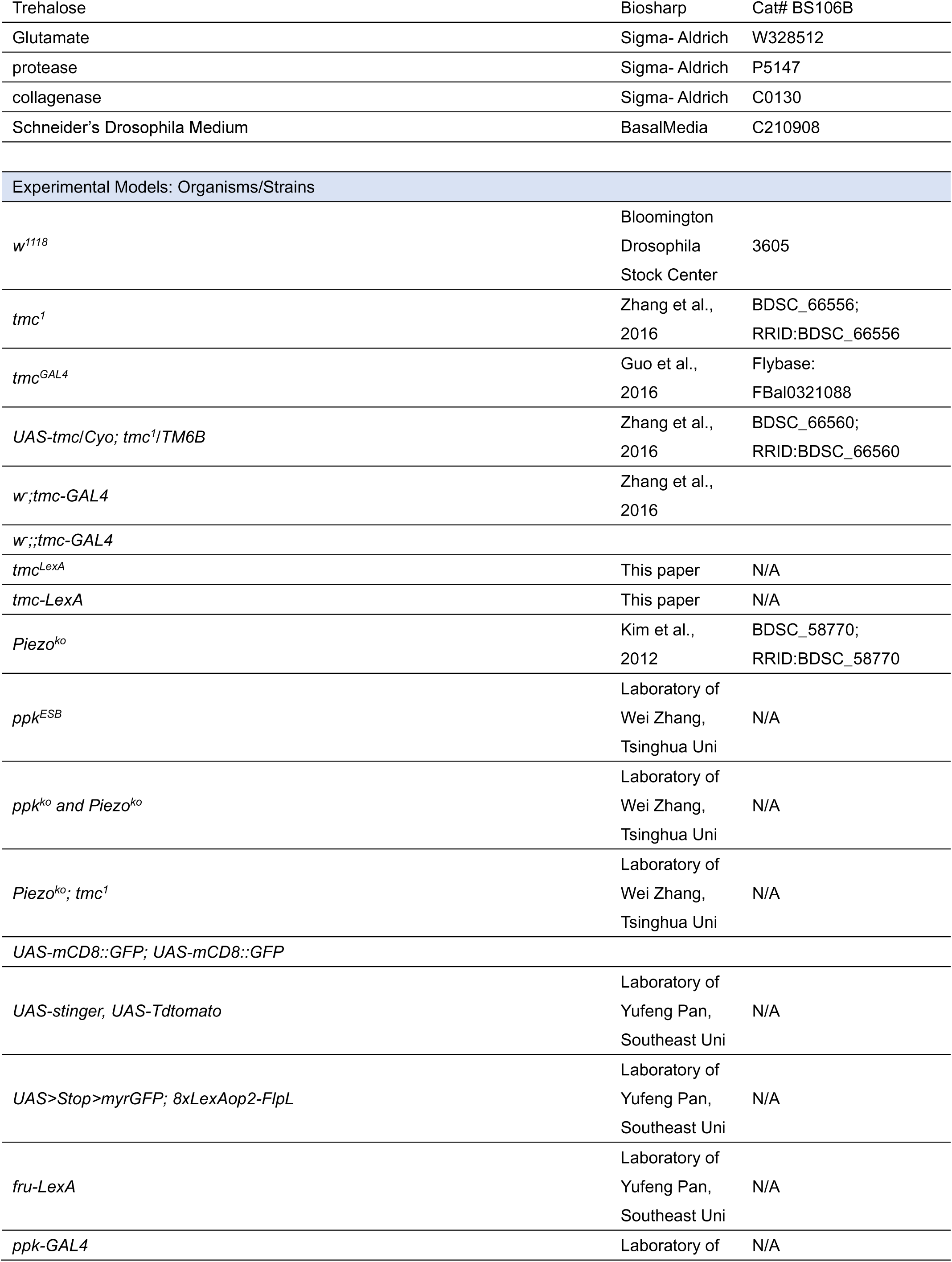
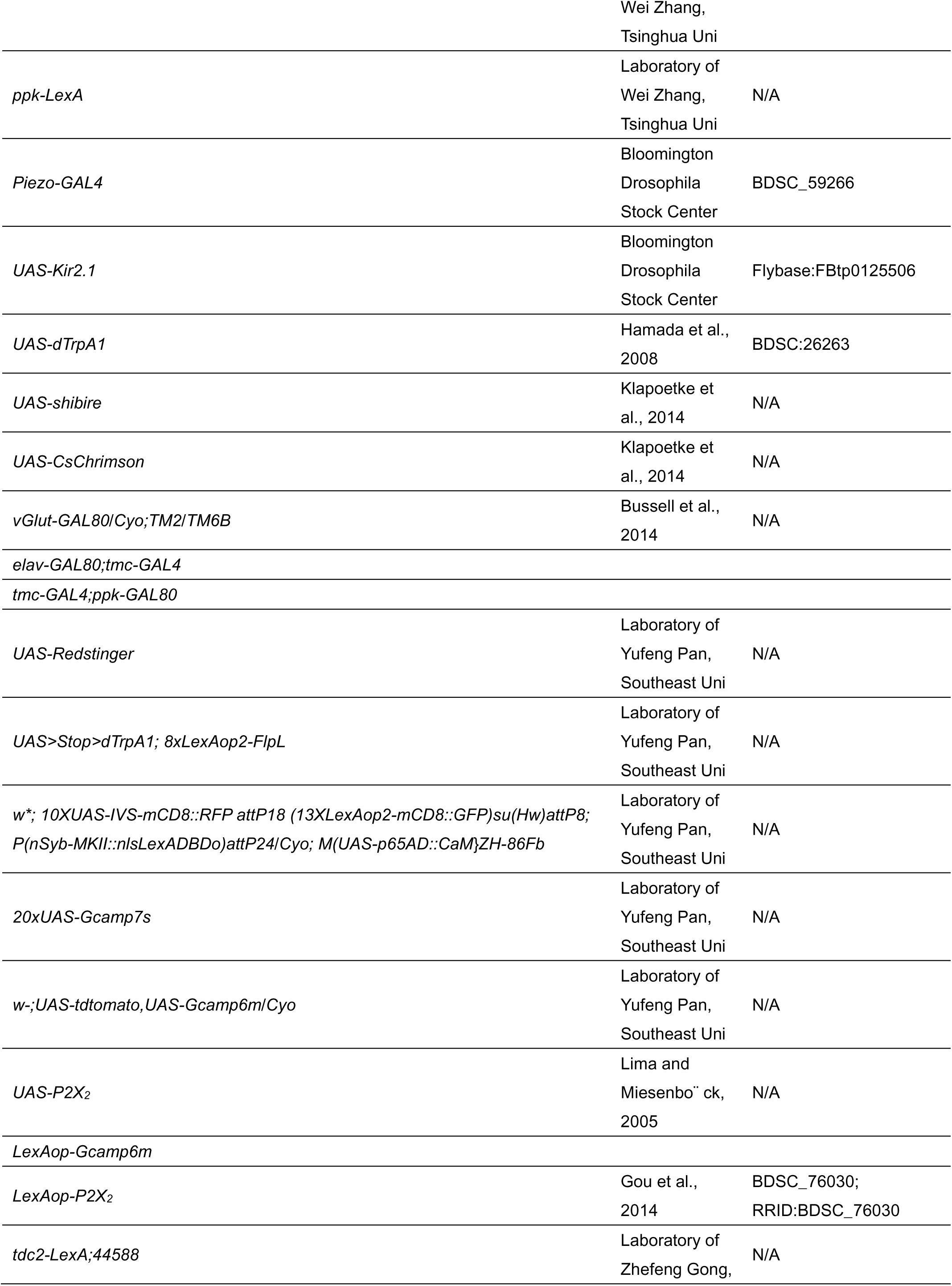
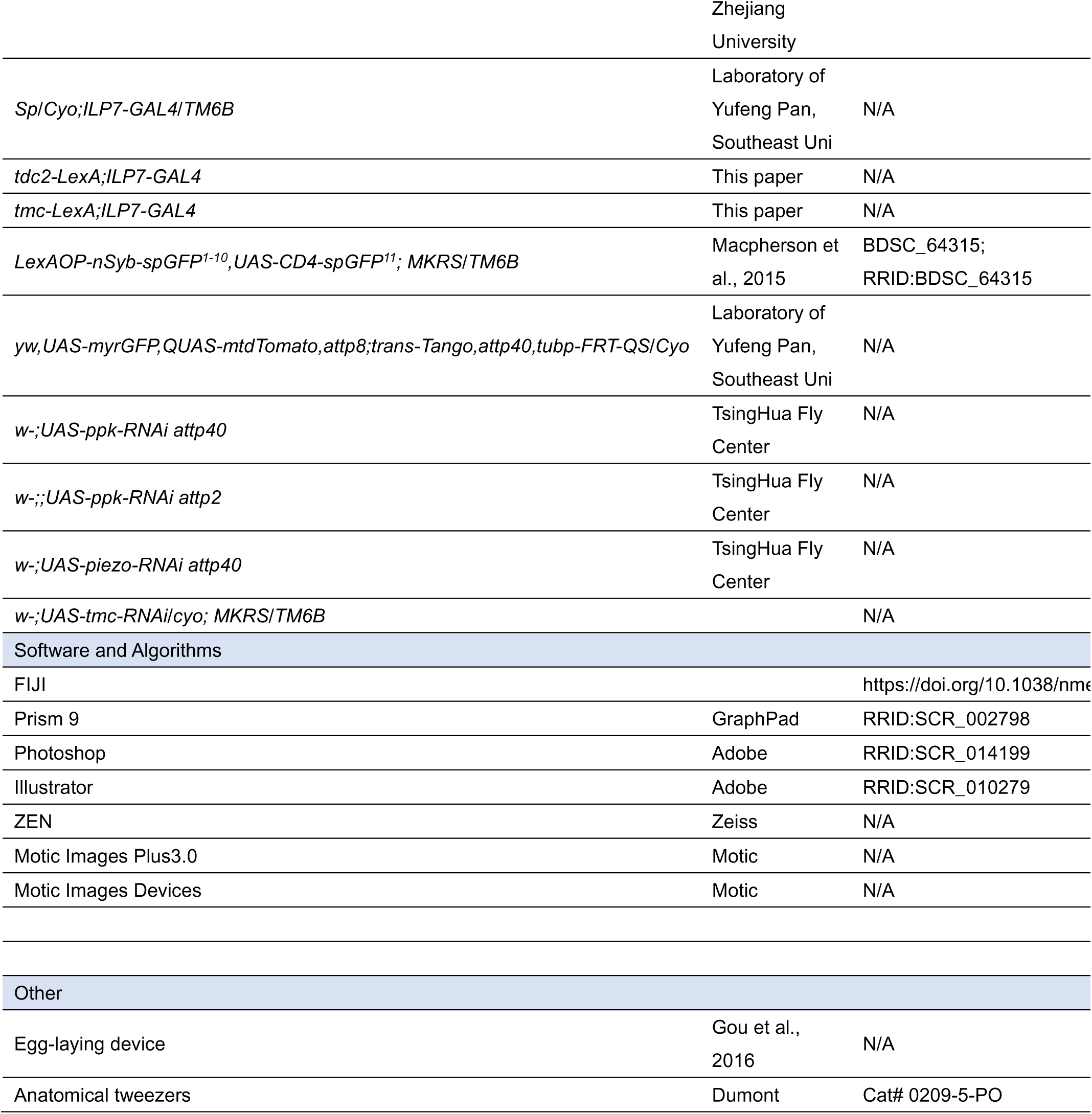
Key resources table.

### Genomic DNA and cDNA amplifications

Genomic DNA was extracted using a buffer (0.1 M Tris-HCl, pH 9.0, 0.1 M EDTA; 1% SDS). In brief, 30 fruit flies were ground with steel beads in the extraction buffer and incubated at 65°C for 1 hour. The samples were then lysed by adding potassium acetate to a final concentration of 800 mM and incubated on ice for 2 hours. Precipitation was pelleted by centrifugation at 10,000g for 15 min, and the supernatant was collected. The same volume of ethanol was added to the supernatant at –20℃ for 2h. Precipitation was pelleted by centrifugation at 10,000g for 15 min, and the sediment was collected. The pellet was washed with ice-cold 70% ethanol once, and the pellet was dried before resuspended in DEPC water.

Total RNA Extraction was done using the TRizol reagent (Invitrogen, Carlsbad, CA, USA) according to the manufacturer’s instructions. The cDNA template used for cloning was synthesized using the Biotech M-MLV reverse transcription kit and the synthesized cDNA template was stored at –20°C

### Egg-laying assays

Unless otherwise mentioned, all egg laying assays were performed with females in darkness with temperature and humidity controlled at 25°C and 60%, respectively.

The experimental flies were prepared as previously described (Gou et al., 2016; Yang et al., 2015; Gou et al., 2014). Briefly, 20 to 30 females of the appropriate genotypes and 10 to 15 males of mixed genotypes were gathered into a single food vial that was supplied with wet yeast paste. After 5-7 days, the food in the vial became very chewed up by the larvae, and at this point, the females were well-fed but deprived of egg-laying. Thus, they were ready to lay eggs when placed in our egg-laying apparatus (Yang et al., 2015; Gou et al., 2016). We usually let females lay eggs overnight (∼14 hr). Groups to be compared were always run in parallel. Note that regardless of the assays we performed, we always assayed at the level of single animals – each data point on a graph denotes the outcome (number of eggs laid) of a single female.

### Assessing ovulation efficiency and egg-jamming ratio

We collected some females for ovary dissection before the egg-laying females were put in the oviposition apparatus, and the remaining females had their ovaries dissected just after the egg-laying experiment in the oviposition apparatus.

We assessed whether there is an ovulation defect by counting mature eggs inside the ovaries of the females before and after the egg-laying assay The ovulation efficiency = (Retained mature eggs before egg-laying – Retained mature eggs after egg-laying) / Retained mature eggs before egg-laying.

Before dissection, we used carbon dioxide for brief anesthesia and severed the connection between the VNC and the brain in the flies. This procedure facilitates dissection and removes the brain’s inhibitory effect on ovulation, allowing for autonomous ovulation behavior in the flies, which aids in subsequent dissection and frequency statistics of egg jamming.

The frequency of egg jamming is determined by dissecting female fruit fly ovaries and noting the position of the eggs in the oviduct. We define egg jamming as the presence of two eggs retained in the lateral oviducts. This definition includes the egg-jamming phenotype caused by the inactivation/activation of *ILP7*-expressing neurons, which also leads to an extra egg being retained in the main oviduct.

### Ovary development measurement

We dissected and made images of the ovaries of virgin flies on d2AAE and d7AAE, respectively. Using Motic Images Devices and Motic Images Plus 3.0 to capture images of the ovaries. By counting the number of retained mature eggs and using FIJI to calculate the ovary’s area and the mature egg’s length and width, we could determine whether the oogenesis and ovary development process was aberrant.

### dTrpA1 activation and shibire inactivation

Experimental flies were maintained at 22°C, cold anesthetized, and allowed to recover for at least 30 min at 22°C. The egg-laying apparatus containing flies was then placed at experimental temperatures (30°C or 22°C) for 14 hours to record the egg-laying numbers.

### CsChrimson activation

For optogenetic stimulation, test flies were collected within 1-3 d after eclosion and transferred into a vial that was supplied with wet yeast paste containing 0.2 mM all-trans-retinal (116–31–4, Sigma-Aldrich). The vials were covered by aluminum foil to protect them from light for 3–4 days before the ovulation. As previously mentioned, we placed the flies in a transparent oviposition device for egg-laying and stimulated the animals with red light (620 nm, 0.03 mW/mm², Vanch Technology, Shanghai, China). Unless otherwise noted, light stimulation was presented continuously throughout the observation period. Light intensity was measured by placing an optical power meter (PS-310 V2, Gentec, Canada) near the location of the glass slide.

### Immunostaining and confocal imaging

We generally dissected the brain, VNC, and ovary of four– to eight-day-old females, except for trans-Tango experiments, in which flies were reared at 20℃ on days 15-20. Dissection was in PBS buffer, and tissues were fixed in 4% PFA solution for ∼25 min at room temperature. Samples were then washed four times for 10 minutes each with PAT3 wash buffer (0.5% Triton X-100, 0.5% bovine serum albumin in PBS). The tissues were transferred to a block buffer (ReadyProbes™ 2.5% Normal Goat Serum) and incubated at room temperature for 60 minutes. Primary antibodies were added to the samples and incubated for 4h at room temperature. Samples were then incubated overnight at 4°C. Samples were washed four times for 10 minutes each and then incubated with secondary antibodies at room temperature for 4h. Samples were washed four times, each for 10 minutes. If DAPI staining was required, the samples were incubated at room temperature with DAPI (1:1000, Solarbio, Cat# C0060) for 30 minutes. Finally, samples were mounted using an anti-fade mounting medium (Vector Laboratories).

Titers for primary antibodies were as follows: mouse anti-nc82 (Developmental Studies Hybridoma Bank nc82, 1:1000), mouse anti-GFP (1:1000, A11120, Thermo Fisher Scientific), rabbit anti-GFP (1:1000, A11122, Thermo Fisher Scientific), and rabbit anti-RFP (1:1000, 600-401-379, Rockland), rabbit anti-TMC (1:250). Secondary antibodies were Donkey Alexa 488 anti-rabbit, Donkey Alexa 488 anti-mouse, Donkey Alexa 555 anti-rabbit, and Donkey Alexa 555 anti-mouse (1:500, R37118, R37114, R37119, R37115, Thermo Fisher Scientific).

Samples were imaged at 10X, 20X, or 40X magnification and 1,024 × 1,024 resolution or more on Zeiss LSM980 confocal microscopes (Jena, Germany) at Nanjing Agricultural University. Images were processed with FIJI or ZEN.

For GRASP, we did not stain the samples with anti-GFP and imaged the fixed and washed samples directly.

For Tdtomato and Stinger, we did not stain the samples with anti-GFP and imaged the fixed and washed samples directly.

For oversized images (such as Fig2J, Fig2S3), we used the Fiji plugin: Stitching-Pixelwise Stitching of Images to process the image stitching.

### Antibody generation

We selected the last 24 amino acids of the TMC protein (C-DPRSASPEPTVNIIRIDIENEHEK) as the peptide antigen for peptide synthesis and conjugated it to a KLH carrier for immunization. The synthesized peptide antigen was injected into rabbits to generate antibodies (ABclonal antibodies). Drosophila TMC protein antibodies were obtained by affinity purification of the antiserum, and the specificity of the antibodies was verified via Dot-Blot(DB) testing against the naked peptide before experiments. For immunolabeling, the TMC antibody was diluted at a 1:250 ratio.

### TRIC

We collected virgin flies with the genotype w*,10XUAS-IVS-mCD8::RFP attP18, {13XLexAop2-mCD8::GFP}su(Hw)attP8; P{nSyb-MKII::nlsLexADBDo}attP24/tmc-GAL4; M{UAS-p65AD::CaM}ZH-86Fb/+ that were enclosed for 5 days and dissected them, separating those that laid eggs and those that did not lay any eggs (number of eggs = 0). After dissection, without staining, we immediately observed the TRIC signal of mdn-LO neurons in real time under the same parameters. We used GFP/RFP in each cell body as the TRIC signal.

### Calcium imaging

For in vitro experiments, VNC and ovaries were dissected in *Drosophila* Adult Hemolymph-Like Saline (AHLS) buffer. The cuticle and connective tissue covering the VNC and ovary were removed using fine forceps. The median abdominal nerve, connecting the abdominal neuromere and the reproductive oviduct, was either kept intact or carefully severed, depending on the experiment.

Calcium imaging was performed on a Zeiss 980 confocal microscope (Jena, Germany) equipped with 10× or 20× objectives. Briefly, cells were scanned with 488 nm and 555 nm lasers at 512 × 512 pixels (or lower) and an 8-bit dynamic range to acquire baseline GCaMP or tdTomato fluorescence. Regions of interest (ROIs) were selected in FIJI, and fluorescence changes were quantified as ΔF/F₀ = (Fₚₑₐₖ – F₀)/F₀, where F₀ is the mean fluorescence over 30 frames in mdn-LO neurons expressing GCaMP. For *ilp7* or *tdc2* neurons, F₀ was defined as the average fluorescence over 30 frames before solution application. Depending on the experiment, solutions were applied as follows: 10 µL AHLS buffer,10 µL of 100 mM ATP, and 10 µL of 1 mM octopamine in 90 µL AHLS; and 20 µL of 50 mM glutamate in 80 µL AHLS.

For positioning the egg within the oviduct, we used an ex vivo “brain-less” preparation. Using forceps, we rapidly displaced the egg to different locations along the oviduct and, after each displacement, promptly refocused and began recording one or more bouts (with brief pauses between bouts to restart acquisition). This allowed us to capture how mechanosensory feedback arising from changes in egg position modulates the activity of ILP7 neurons.

### Single-nucleus RNA-sequencing analyses in *ilp7* neurons

We queried the Fly Cell Atlas (FCA, https://flycellatlas.org/) and focused on the Body (10×, Stringent) dataset of *Drosophila melanogaster* (ASAP). We performed a three-gene co-expression screen: (1) *nSyb* was used as a neuronal marker to restrict the analysis to VNC neurons; (2) *Ilp7* was used to identify *Ilp7*-positive cells; and (3) each candidate gene from our curated list was queried within that set. Cells co-expressing *nSyb* and *Ilp7* were operationally defined as *Ilp7* neurons. For each candidate gene, we recorded its expression within these Ilp7 neurons and mapped the common co-expression to five FCA cell coordinate codes: (−13.14, 0.14), (−13.34, 0.24), (−14.19, −1.32), (−15.20, −0.17), (−15.37, −1.57). Gene expression levels are reported as Norm_1_asap_seurat values as provided by the FCA portal.

### Quantification and statistical analysis

We used the GraphPad Prism6 software package to generate graphs and analyze data statistically. We tested whether the values were normally distributed using D’Agostino–Pearson omnibus and Shapiro–Wilk normality tests before performing statistical analysis. When data were normally distributed, we used parametric tests; when data were not normally distributed, we used non-parametric tests. All tests were two-tailed. All data are presented as mean ± s.e.m. We also consistently labeled the sample numbers directly on graphs.

### Sample size

Our results were based on an analysis of behavior at the level of single animals.

Sample sizes were determined before experimentation based on both variance and effect sizes. All the data are biological replicates.

## Acknowledgments

This research was funded by the the National Natural Science Foundation of China (32022011 & 32472542), the National Key Research and Developmental Program of China (2022YFD1700200) and the Fundamental Research Funds for Central Universities of the Central South University (KJJQ2025014).

## Movie legend

Movie S1. The video shows that during lateral oviduct contractions, Ca²⁺ levels in mdn-LO neurons increase in phase with muscle contraction, indicating responsiveness to lateral oviduct activity.

Movie S2. With the median abdominal nerve (MAN) intact, Ca²⁺ in mdn-LO axons within the VNC region of the VNC rises markedly during lateral oviduct contractions.

Movie S3. Following transection of the MAN, Ca²⁺ in mdn-LO axons within the VNC region shows no significant increase during lateral oviduct contractions, indicating that mdn-LO conveys signals to the VNC via the MAN.

Movie S4. ATP stimulation of P2X2-expressing mdn-LO neurons produces a pronounced elevation of Ca²⁺ in *ilp7* neurons within the VNC and triggers strong contractions of the lateral oviduct musculature.

Movie S5. The oscillation of the female-specific post-embryonic ILP7 neurons.

Movie S6. The oscillation of the embryonic ILP7 neurons.

Movie S7. The oscillation of the dissociated ILP7 neurons.

